# Spatiotemporal coding in the macaque supplementary eye fields: landmark influence in the target-to-gaze transformation

**DOI:** 10.1101/2020.06.25.172031

**Authors:** Vishal Bharmauria, Amirsaman Sajad, Xiaogang Yan, Hongying Wang, John Douglas Crawford

**Author notes:** **Corresponding author:** Dr. John Douglas Crawford, Departments of Psychology, Biology and Kinesiology & Health Sciences, York University, Toronto, Canada, Centre for Vision Research, Room 0009A LAS, 4700 Keele Street, Toronto, Ontario, M3J 1P3. **Significance Statement** It is thought that the brain integrates egocentric (self-centered) and allocentric (landmark-centered) visual signals to generate accurate goal-directed movements, but the neural mechanism is not known. Here, by shifting a visual landmark while recording frontal cortex activity in awake behaving monkeys, we show that the supplementary eye fields (SEF) incorporates landmark-centered information (in memory and motor activity) when it transforms target location into future gaze position commands. We propose a circuit model in which the SEF provides control signals to implement an integrated gaze command in the frontal eye fields (Bharmauria et al., 2020). Taken together, these experiments explain normal ego / allocentric integration and might suggest rehabilitation strategies for neurological patients who have lost one of these visual mechanisms.

## Abstract

Eye-centered (egocentric) and landmark-centered (allocentric) visual signals influence spatial cognition, navigation and goal-directed action, but the neural mechanisms that integrate these signals for motor control are poorly understood. A likely candidate for ego / allocentric integration in the gaze control system is the supplementary eye fields (SEF), a mediofrontal structure with high-level ‘executive’ functions, spatially tuned visual / motor response fields, and reciprocal projections with the frontal eye fields (FEF). To test this hypothesis, we trained two head-unrestrained animals to saccade toward a remembered visual target in the presence of a visual landmark that shifted during the delay, causing gaze end points to shift partially in the same direction. 256 SEF neurons were recorded, including 68 with spatially tuned response fields. Model fits to the latter established that, like the FEF and superior colliculus, spatially tuned SEF responses primarily showed an egocentric (eye-centered) target-to-gaze position transformation. However, the landmark shift influenced this default egocentric transformation: during the delay, motor neurons (with no visual response) showed a transient but unintegrated shift (i.e., not correlated with the target-to-gaze transformation), whereas during the saccade-related burst visuomotor neurons showed an integrated shift (i.e., correlated with the target-to-gaze transformation). This differed from our simultaneous FEF recordings (Bharmauria et al., 2020), which showed a transient shift in visuomotor neurons, followed by an integrated response in all motor responses. Based on these findings and past literature, we propose that prefrontal cortex incorporates landmark-centered information into a distributed, eye-centered target-to-gaze transformation through a reciprocal prefrontal circuit.

## INTRODUCTION

The brain integrates egocentric (eye-centered) and allocentric (landmark-centered) visual cues to guide goal-directed behavior (Ball et al., 2009; Chen et al., 2011; Goodale and Haffenden, 1998; Karimpur et al., 2020). For example, to score a goal, a soccer forward must derive allocentric relationships (e.g. where is the goaltender relative to the posts?) from eye-centered visual inputs to predict an opening, and then transform this into body-centered motor commands. The neural mechanisms for ego / allocentric coding for vision, cognition, and navigation have been extensively studied (Ekstrom et al., 2014; Milner and Goodale, 2006; O’Keefe, 1976; O’Keefe and Dostrovsky, 1971; Rosenbaum et al., 2004; Schenk, 2006), but the mechanisms for goal-directed behavior are poorly understood. One clue is that humans aim reaches toward some intermediate point between conflicting ego / allocentric cues, suggesting Bayesian integration (Bridgeman et al., 1997; Byrne and Crawford, 2010; Fiehler et al., 2014; Klinghammer et al., 2017; Lemay et al., 2004; Neely et al., 2008). Neuroimaging studies suggest this may occur in parietofrontal cortex (Chen et al., 2018) but could not reveal the cellular mechanisms. However, similar behavior has been observed in the primate gaze system (Li et al., 2017), suggesting this system can be also used to study ego / allocentric integration.

It is thought that higher level gaze structures—lateral intraparietal cortex (LIP), frontal eye fields (FEF), superior colliculus (SC)— primarily employ eye-centered codes (Goldberg et al., 2002; Klier et al., 2001; Paré and Wurtz, 2001; Russo and Bruce, 1993; Tehovnik et al., 2000), and transform target location (T) into future gaze position (G) (Constantin et al., 2007; Everling et al., 1999; Schall et al., 1995). We recently confirmed this by fitting various spatial models against FEF and SC response field activity (Sadeh et al., 2020, 2015; Sajad et al., 2016, 2015). Visual responses coded for target position in eye coordinates (Te), whereas motor responses (separated from vision by a delay) coded for future gaze position in eye coordinates (Ge), and a progressive target-to-gaze (T-G) transformation (where G includes errors relative to T) along visual-memory-motor activity. Further, when we introduced a visual landmark, and then shifted it during the delay period, response field coordinate systems (in cells with visual, delay, and motor activity) shifted partially in the same direction (Bharmauria et al., 2020). Eventually this allocentric shift became integrated into the egocentric (T-G) transformation of all cells that produced a motor burst during the saccade. However, it is unclear if this occurs independently through a direct visuomotor path to FEF, or in concert with higher control mechanisms.

One likely executive control mechanism could be the supplementary eye fields (SEF), located in the dorsomedial frontal cortex (Schlag and Schlag-Rey, 1987) and reciprocally connected to the FEF (Huerta and Kaas, 1990; Stuphorn, 2015). The SEF has visual and gaze motor response fields (Schall, 1991; Schlag and Schlag-Rey, 1987), but its role in gaze control is controversial (Abzug and Sommer, 2017). The SEF is involved in various high-level oculomotor functions (Olson and Gettner, 1995; Sajad et al., 2019; Stuphorn et al., 2000; Tremblay et al., 2002) and reference frame transformations, both egocentric (Martinez-Trujillo et al., 2004; Schall et al., 1993; Schlag and Schlag-Rey, 1987), and object-centered, i.e., one part of an object relative to another (Olson and Gettner, 1995; Tremblay et al., 2002). However, the role of the SEF in coding egocentric visuomotor signals for head-unrestrained gaze shifts is untested, and its role in the implicit coding of gaze relative to independent visual landmarks is unknown.

Here, we recorded SEF neurons in two head-unrestrained monkeys while they performed gaze shifts in the presence of implicitly conflicting egocentric and allocentric cues (**Fig. 1A**). As reported previously, gaze shifted away from the target, in the direction of a shifted landmark (Byrne and Crawford, 2010; Li et al., 2017). We employed our model-fitting procedure (Keith et al. 2009; DeSouza et al. 2011; Sadeh et al. 2015; Sajad et al. 2015) to analyze the neural data. First, we tested all possible egocentric and allocentric models. Second, we performed a spatial continuum analysis (between both egocentric and allocentric models) through time to see if, when, and how egocentric and allocentric transformations are integrated in the SEF. We find that 1) SEF neurons predominantly possess an eye-centered transformation from target-to-gaze coding and 2) landmark-centered information is integrated into this transformation, but through somewhat different cellular mechanisms than the FEF (Bharmauria et al., 2020). We thus propose a reciprocal SEF-FEF model for allo / egocentric integration in the gaze system.

**Figure 1.**
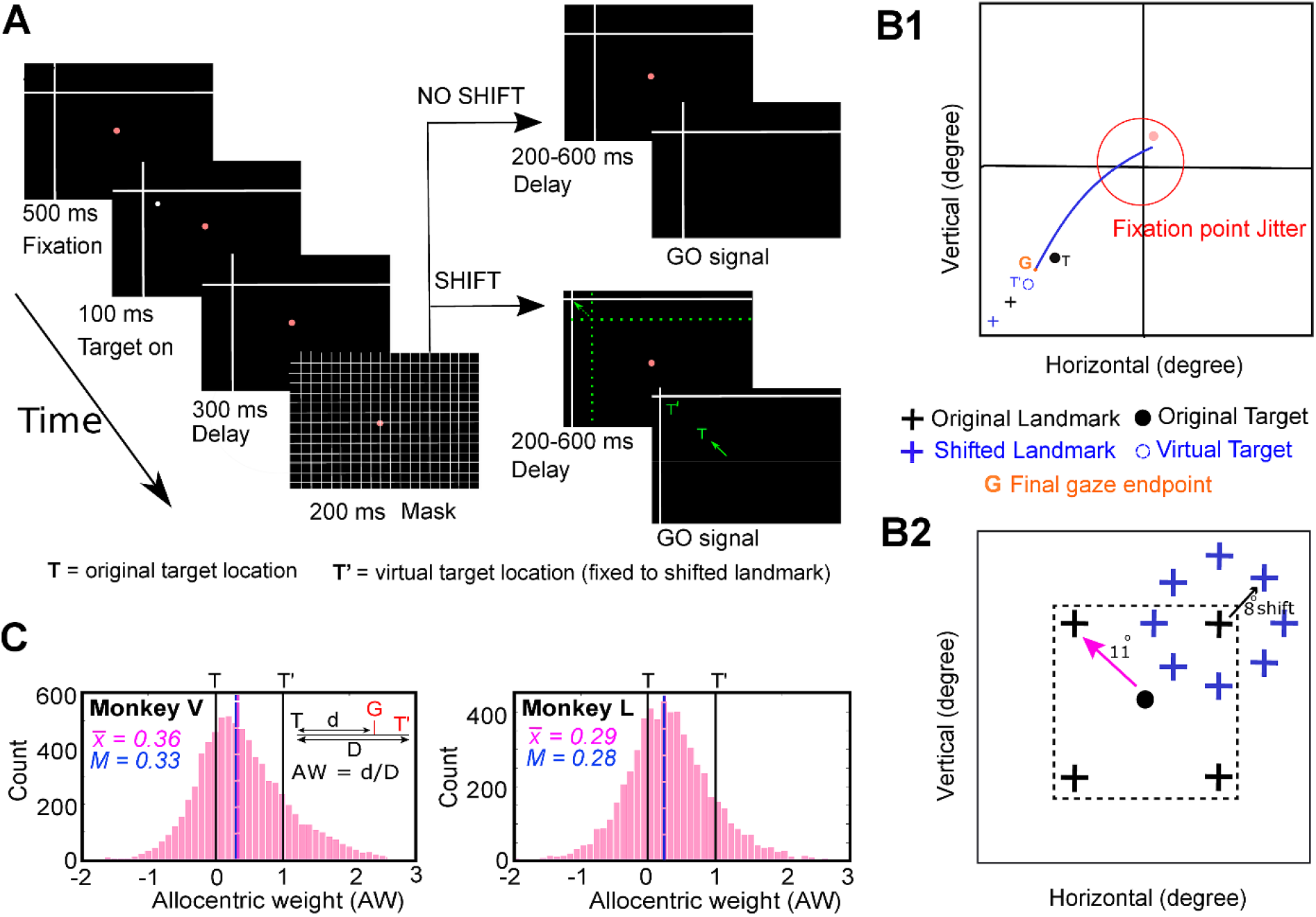
Experimental paradigm and behavior. **(A)** Cue-conflict paradigm and its timeline. The monkey began the trial by fixating for 500 ms on a red dot in the presence of a landmark (L, intersecting lines) was already present on the screen. Then a target (white dot) was presented for 100 ms, followed by 300 ms delay and a grid-like mask (200 ms). After the mask, and a second memory delay (200-600 ms), the animals was signaled (extinguishing of the fixation dot, i.e., go signal) to saccade head-unrestrained to the remembered target location either when the landmark was shifted (L’, denoted by the broken green arrow) or when it was not shifted (i.e., present at the original location). The monkey was rewarded for landing his gaze (G) within a radius of 8-12° around the original target (monkeys were rewarded for looking at T = original target, at T’ = virtually shifted target fixed to landmark, or between T and T’). The green arrow represents the head-unrestrained gaze shift to the remembered location. The reward window was centered on T so that the behavior was not biased. Note that the actual landmark shift was 8°, but for clarity, it has been increased in this schematic figure as indicated by the broken green arrow. Notably, the green colored items are only for the purposes of representaion (they were not present on the screen). **(B1)** Schematic of a gaze shift (blue line) from the home fixation point (red dot) toward the virtual target (broken blue circle, T’) fixed to the shifted landmark (blue cross, L’). ‘G’ refers to the final gaze endpoint. The blue square represents the area in B2. **(B2)** Schematic illustration of all (four) possible locations of the landmark (black cross) for an example target (black dot, T), with all possible locations of the shifted landmark relative to original landmark location. The landmark was presented 11° (indicated by pink arrow) from the target, in one of the four oblique directions and post-mask it shifted 8° (blue cross stands for the shifted landmark, black arrow depicts the shift) from its initial location in one of the eight radial directions around the original landmark **. C)** Allocentric weight (AW) distribution (X-axis) for monkey V (left) and monkey L (right) plotted as a function of the number of trials (Y-axis) for all the analyzed trials from the spatially tuned neurons. The mean AW (vertical pink line) for MV and ML was 0.36 and 0.29 (mean = 0.33) respectively. **Note:** the ‘shift’ condition included 90 % of trials and the ‘no shift’ condition only included 10 % of the trials. The AW was calculated for the shifted condition only. The ‘no-shift’ condition allowed to test for all the range of shifts. To not bias the behavior, the reward window included scores from – 1 - + 1.

## MATERIALS AND METHODS

### Surgical Procedures and Recordings of 3D Gaze, Eye, and Head

The experimental protocols complied the guidelines of Canadian Council on Animal Care on the use of laboratory animals and were approved by the York University Animal Care Committee. Neural recordings were performed on two female *Macaca mulatta* monkeys (Monkey V and Monkey L) and they were implanted with 2D and 3D sclera search coils in left eyes for eye-movement and electrophysiological recordings (Crawford et al., 1999; Klier et al., 2003). The eye coils allowed us to register 3D eye movements (i.e., gaze) and orientation (horizontal, vertical, and torsional components of eye orientation relative to space). During the experiment, two head coils (orthogonal to each other) were also mounted that allowed similar recordings of the head orientation in space. We then implanted the recording chamber centered in stereotaxic coordinates at 25 mm anterior and 0 mm lateral for both animals. A craniotomy of 19 mm (diameter) that covered the chamber base (adhered over the trephination with dental acrylic) allowed access to the right SEF. Animals were seated within a custom-made primate chair during experiments, and this allowed free head movements at the center of three mutually orthogonal magnetic fields (Crawford et al., 1999). The values acquired from 2-D and 3-D eye and head coils allowed us to compute other variables such as the orientation of the eye relative to the head, eye- and head-velocities, and accelerations (Crawford et al., 1999).

### Basic Behavioral Paradigm

The visual stimuli were presented on a flat screen (placed 80 cm away from the animal) using laser projections **(Fig. 1A).** The monkeys were trained on a standard memory-guided gaze task to remember a target relative to a visual allocentric landmark (two intersecting lines acted as an allocentric landmark) thus leading to a temporal delay between the target presentation and initiation of the eye movement. This allowed us to separately analyze visual (aligned to target) and eye-movement-related (saccade onset) responses in the SEF. To not provide any additional allocentric cues, the experiment was conducted in a dark room. In a single trial, the animal began by fixating on a red dot (placed centrally) for 500 ms while the landmark was present on the screen. This was followed by a brief flash of visual target (T, white dot) for 100 ms, and then a brief delay (300 ms), a grid-like mask (200 ms, this hides the past visual traces, and also the current and future landmark) and a second memory delay (200-600 ms, i.e., from the onset of the landmark until the go signal). As the red fixation dot extinguished, the animal was signaled to saccade head-unrestrained (indicated by the solid green arrow) toward the memorized location of the target either in the presence of a shifted (indicated by broken green arrow) landmark (90 % of trials) or in absence of it (10 %, no-shift/zero-shift condition, i.e. landmark was present at the same location as before mask). These trials with zero-shift were used to compute data at the ‘origin’ of the coordinate system for the T-T’ spatial model fits as described below. The saccade targets were flashed one-by-one randomly throughout the response field of a neuron. Note: green color highlights the items that were not presented on the screen (they are only for representational purposes).

The spatial details of the task are in **Figure 1B**. **Figure 1B1** shows an illustration of a gaze shift (blue curve) to an example target (T) in presence of a shifted landmark (blue cross). ‘G’ refers to the final gaze endpoint and T’ stands for the virtual target (fixed to the shifted landmark). The landmark vertex could initially appear at one of four spots located 11° obliquely relative to the target and then shift in any one of 8 directions (**B2**). Importantly, *the timing and amplitude (8°) of this shift was fixed*. Since these animals had been trained, tested behaviorally (Li et al., 2017) and then retrained for this study over a period exceeding two years, it is reasonable to expect that they may have learned to anticipate the timing and the amount of influence of the landmark shift. However, we were careful not to bias this influence: animals were rewarded with a water-drop if gaze was placed (G) within 8-12° radius around the original target (i.e., they were rewarded if they looked at T, toward or away from T’, or anywhere in between). Based on our previous behavioral result in these animals (Li et al., 2017), we expected this paradigm to cause gaze to shift partially toward the virtually shifted target in landmark coordinates (T’).

Note that this paradigm was optimized for our method for fitting spatial models to neural activity (see below), which is based on variable dissociations between measurable parameters such as target location and effectors (gaze, eye, head), and various egocentric / allocentric reference frames (Keith et al., 2009; Sajad et al., 2015). This was optimized by providing variable landmark locations and shift directions, and the use of a large reward window to allow these shifts (and other endogenous factors) to influence gaze errors relative to T. We also jittered the initial fixation locations within a 7-12° window to dissociate gaze-centered and space-centered frames of reference (note that no correlation was observed between the initial gaze location and final gaze errors). Further dissociations between effectors and egocentric frames were provided by the animals themselves, i.e., in the naturally variable contributions of the eye and head to initial gaze position and the amplitude/direction of gaze shifts. Details of such behavior have been described in detail in our previous papers (e.g., Sadeh et al., 2015; Sajad et al., 2015).

### Behavioral Recordings and Analysis

During experiments, we recorded the movement of eye and head orientations (in space) with a sampling rate of 1000 Hz. For the analysis of eye movement, the saccade onset (eye movement in space) was marked at the point in time when the gaze velocity exceeded 50°/s and the gaze offset was marked as the point in time when the velocity declined below 30°/s. The head movement was marked from the saccade onset till the time point at which the head velocity declined below 15°/s.

When the landmark shifted (90% of trials), its influence on measured future gaze position (G) was called allocentric weight (AW), computed as follows:

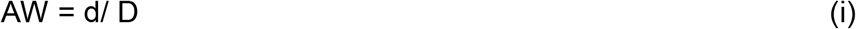

where *AW* is allocentric weight; *d* is the component of T-G (error space between the actual target location and the final measured gaze position) that projects onto the vector direction of the landmark shift, and D is the magnitude of the landmark shift (Byrne and Crawford, 2010; Li et al., 2017). This was done for each trial, and then averaged to find the representative landmark influence on behavior in a large number of trials. A mean AW score of zero signifies no landmark influence, i.e., gaze shifts headed on average toward T. A mean AW score of 1.0 means that on average, gaze headed toward a virtual target position (T’) that remained fixed to the shifted landmark position. As we shall see (**Fig. 1 C**), AW scores for individual trials often fell between 0 and 1 but varied considerably, possibly due to trial-to-trial variations in landmark influence and / or other sources of variable gaze error that are present without a landmark shift (Sajad et al., 2020, 2016, 2015).

### Electrophysiological Recordings and Response Field Mapping

Tungsten electrodes (0.2–2.0 mΩ impedance, FHC Inc.) were lowered into the SEF [using Narishige (MO-90) hydraulic micromanipulator] to record extracellular activity. Then the recorded activity was digitized, amplified, filtered, and saved for offline spike sorting using template matching and applying principal component analysis on the isolated clusters with Plexon MAP System. The recorded sites of SEF (in head-restrained conditions) were further confirmed by injecting a low-threshold electrical microstimulation (50 μA) as previously used (Bruce et al., 1985). A superimposed pictorial of the recorded sites from both animals is presented in **Figure 2A-B** (Monkey L in Blue and Monkey V in red).

**Figure 2.**
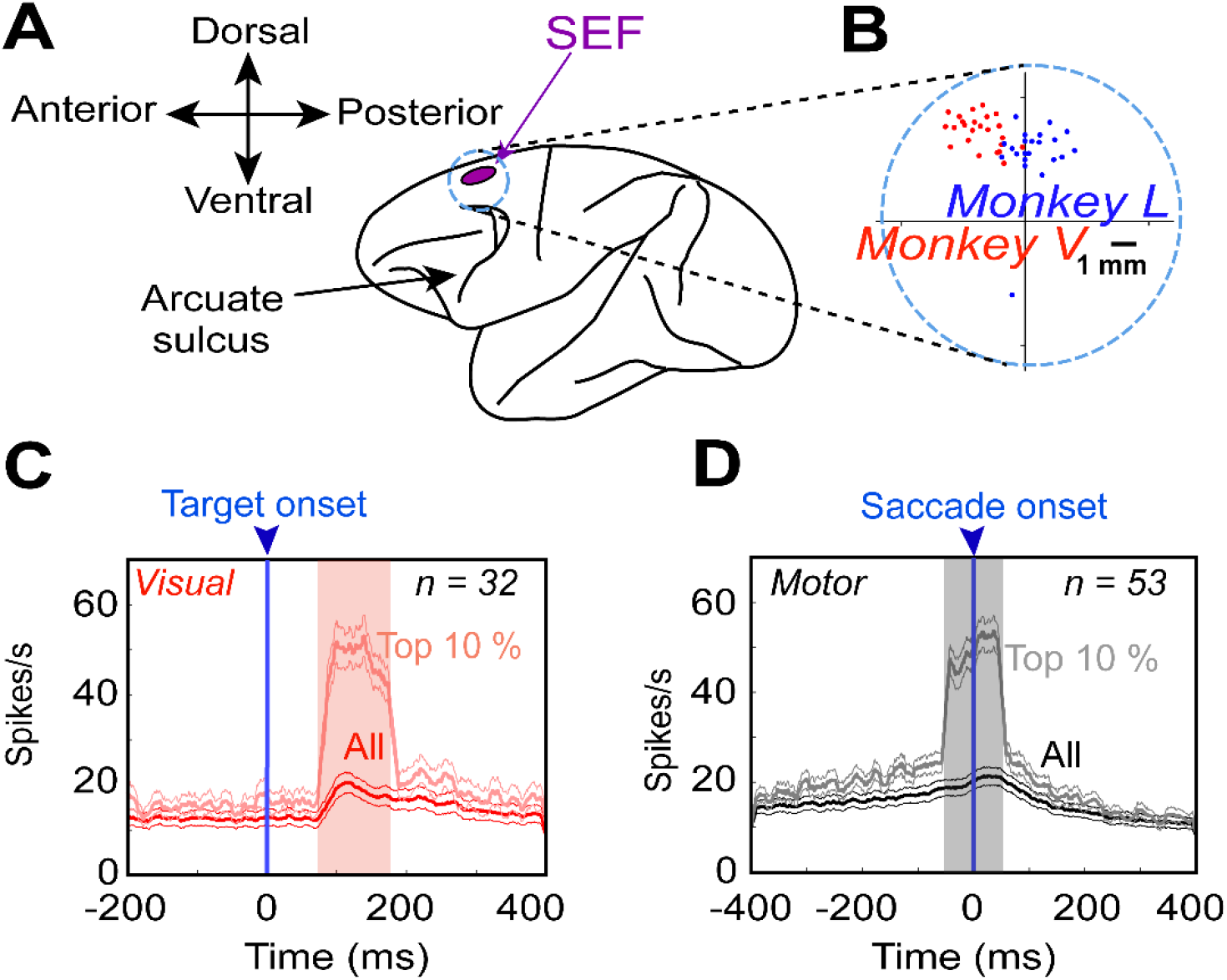
SEF recordings. **(A)** The purple insert represents the location of the SEF and the circle (note that it does not correspond to the original size of chamber) corresponds to the chamber. **(B)** A zoomed-in overlapped section of the chamber and the sites of neural recordings (dots) confirmed with 50 μA current stimulation from Monkey V (red) and Monkey L (blue). **(C)** Mean (± 95 % confidence) of the spike-density plots [dark red: all trials from all neurons; light red: top 10% best trials most likely representing the hot spot of every neuron’s response field in the visual population, aligned to the onset of the target (blue arrow)]. **(D)** Same as **C,** but for motor responses aligned to the saccade onset (blue arrow). The shaded region denotes the 100-ms sampling window. Note that, in both plots, the ‘top 10%’ neural data were selected using the above sampling windows, therefore, delay activity is not completely represented in these plots (it will be examined later in results).

We mostly searched for neurons while the monkey freely (head-unrestrained) scanned the environment. Once we noticed that a neuron had reliable spiking, the experiment started. The response field (visual and/or motor) of neuron was characterized while the animal performed the memory-guided gaze shift paradigm as described above. After an initial sampling period to determine the horizontal and vertical extent of the response field, targets were presented (one per trial) in a 4 × 4 to 7 × 7 array (5 –10° from each other) spanning 30-80°. This allowed the mapping of visual and motor response fields such as those shown in the ‘Results’ Section. For analysis we aimed at approximately 10 trials per target, so the bigger the response field (and thus the more targets), the more the number of recorded trials were required and vice versa. On average 343 ± 166 (mean ± SD) trials/neuron were recorded, again depending on the size of the response field. We did such recordings from > 200 SEF sites, often in conjunction with simultaneous FEF recordings, as reported previously (Bharmauria et al., 2020).

### Data Inclusion Criteria, sampling window and neuronal classification

In total, we isolated 256 SEF neurons: 102 and 154 neurons were recorded from Monkeys V and L respectively. Of these, we only analyzed task-modulated neurons with clear visual burst and/or with perisaccadic movement response **(Fig. 2C-D**). Neurons that only had post-saccadic activity (activity after the saccade onset) were excluded. Moreover, neurons that lacked significant spatial tuning were also eliminated (see ‘Testing for Spatial Tuning’ below). In the end, after applying our exclusion criteria, we were left with 37 and 31 spatially tuned neurons in Monkeys V and L respectively. Only those trials were included where monkeys landed their gaze within the reward acceptance window, however, we eliminated gaze end points beyond ± 2° of the mean distribution from our analysis. For the analysis of the neural activity, the “visual epoch” sampling window was chosen as a fixed 100-ms window of 80 −180 ms aligned to the target onset, and the “movement period” was characterized as a high-frequency perisaccadic 100-ms (−50 - +50 ms relative to saccade onset) period (Sajad et al., 2015). This allowed us to get a good signal/noise ratio for neuronal activity analysis, and most likely corresponded to the epoch in which gaze shifts were influenced by SEF activity. After rigorously employing our exclusion criteria, we further dissociated the spatially tuned neurons into pure visual (V, n =6), pure motor (M, n = 26) and visuomotor neurons (VM — which possessed visual, maximal delay, and motor activity, n = 36) based on the common dissociation procedure (Bruce and Goldberg, 1985; Khanna et al., 2019; Sajad et al., 2015; Schall, 2015). Note: the motor population also includes neurons which have delayed memory activity (Sajad et al., 2016).

### Spatial Models Analyzed (Egocentric and Allocentric)

**Figure 3** graphically shows how we derived the eleven egocentric “canonical” models tested in this study and then chose the best egocentric model to compare with the allocentric models. Briefly, based on these models, this figure illustrates the formal means for comparing between target vs. gaze position coding and gaze vs. eye vs. head displacement/position **(Fig. 3A-C),** with each plotted in different possible frames of reference. We then use the landmark-centered information to test between the egocentric vs. allocentric modes of coding **(Fig. 3D).** Note that in our analysis, these models were based on actual target, gaze, eye, and head data, either derived from geometric calculations (in the case of T) or eye/head coil measurements. As described previously, the variability required to distinguish between these models was either provided by ourselves (i.e., stimulus placement) or the monkeys’ own behavior (i.e., variable gaze errors and variable combinations of eye-head position) (Sajad et al., 2015).

**Figure 3.**
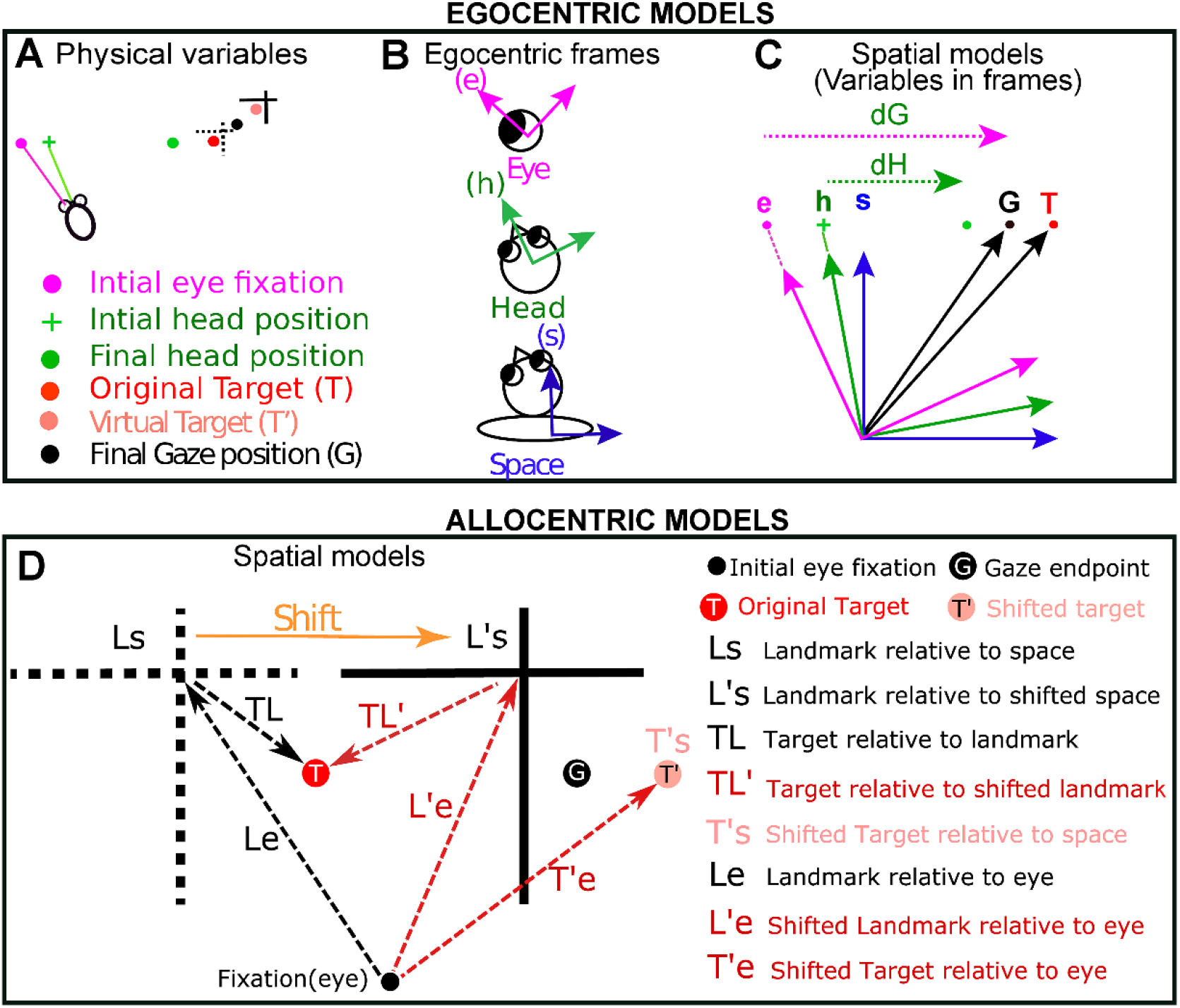
Description of the different spatial models (egocentric and allocentric) tested in this study. **(A)** Physical variables in one trial. The projections of initial gaze (magenta dot), initial head (green cross), final gaze (black dot), and final head (green dot) orientations on the screen are depicted. Note: final gaze often lands between the target (T, red dot) and shifted target (T’, light red dot). **(B)** While head-unrestrained, these positions can be encoded in relation to three egocentric frames of reference: eye (e), head (h), and space/body (s). Since the target was sufficiently far from the head to minimize translation effects, and because our model-fitting method is insensitive to these biases, we centered these frames to be aligned as shown in C. **(C)** Different canonical models derived after plotting the physical parameters in fundamental egocentric frames. dG: gaze displacement referring to final gaze position with respect to the home fixation point (not the eye). dH: displacement of head in space coordinates. dE (not shown) is the displacement of eye in the head accompanying the gaze shift. Target and final gaze, eye, and head positions can be represented in any one of the three initial reference frames. See text for details of all egocentric models **(D)** Allocentric models (see below) that were compared with the most relevant egocentric models (Te, Ge). The broken black cross represents the location of initial landmark (L) and the solid black cross stands for the shifted (indicated by orange arrow) landmark (L’). The red, the black and the light red solid circles correspond to the Target (T), the Gaze endpoint (G) and the virtual Target (T’) locations respectively. Tested Allocentric models: Ls, Landmark relative to space; Le, Landmark relative to eye; TL, Target relative to landmark; L’s, shifted Landmark relative to space; L’e, shifted Landmark relative to eye, TL’, Target relative to shifted landmark, T’e, shifted Target relative to eye; T’s shifted Target relative to space.

We first tested for all the egocentric models as reported in our previous studies (Sajad et al., 2016, 2015) and then tested the best egocentric model with all ‘pure’ allocentric models. Based on different spatial parameters (most importantly Target, T, and final Gaze position, G), we fitted visual and motor response fields of SEF against previously tested eleven **(Fig. 3A-C)** egocentric canonical models in FEF (Sajad et al., 2015). Note that Te and Ge codes were obtained by mathematically rotating the experimental measures of T and G in space coordinates by the inverse of experimentally measured initial eye orientation to obtain the measures in eye coordinates (Klier et al., 2001). In other words, Te is based on the actual target location relative to initial eye orientation, whereas Ge is based upon the final gaze position relative to initial eye orientation.

The following models were also tested: dH, difference between the initial and final head orientation in space coordinates; dE: difference between the initial and final eye orientation in head coordinates; dG, difference between the initial and final gaze position in space coordinates; Hs, final position of the head in space coordinates; Eh, final position of the eye in head coordinates; Gs, Ge, Gh — Gaze in space, eye, and coordinates respectively; Ts, Te, Th — Target in space, eye and head coordinates respectively. The final position refers to orientation of eye/head after the gaze saccade. These models are detailed in the previous study (Sajad et al., 2015). Note that some of these models (like dG and Ge) might not be distinguishable within the range of initial gaze positions and saccade amplitudes described here (Crawford and Guitton, 1997).

Since an allocentric landmark was involved in this task, we analyzed eight additional (allocentric) models of target coding based on the original and shifted landmark location **(Fig. 3D)**. In the allocentric analysis, we retained the best egocentric model (Te for visual neurons and Ge for motor neurons) for comparison. The tested allocentric models are: Ls, Landmark within the space coordinates; L’s, Shifted landmark within the space coordinates; Le, Landmark within eye coordinates; L’e, Shifted landmark within eye coordinates; T’s, Shifted target in space coordinates; T’e, Shifted target in eye coordinates; TL, Target relative to landmark; TL’, target relative to shifted landmark. Note: prime (’) stands for the positions that are related to the shifted location of the landmark.

### Intermediate spatial models used in main analysis

Previous reports on FEF responses from our lab have reported that responses do not fit exactly against spatial models like Te or Ge, but actually may fit best against intermediate models between the canonical ones (**Fig. 4A**, lower left) (Sajad et al., 2020). As in our previous studies (Sadeh et al., 2020; Sajad et al., 2016), we found that a T-G continuum (specifically, steps along the ‘error line’ between Te to Ge) best quantified the SEF egocentric transformation **(Fig. 4A**, lower left). This continuum is similar to the concept of an intermediate frame of reference (e.g., between the eye and head) but is instead intermediate between target and gaze position within the same frame of reference. Hereafter, we will sometimes refer to T-G as ‘the egocentric code’.

**Figure 4.**
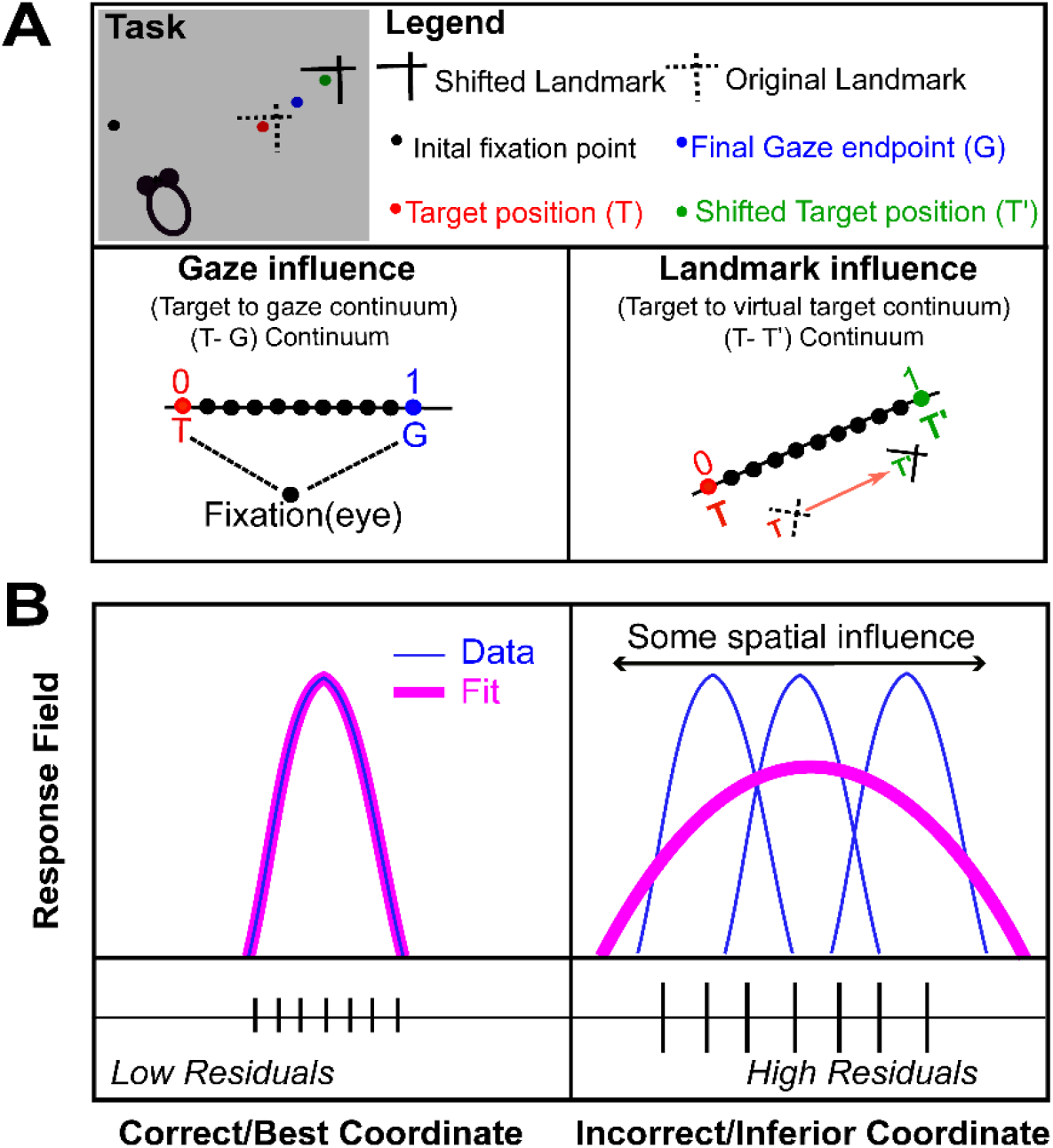
Schematic representation of spatial parameters and spatial model fitting technique. **(A)** An illustration of different spatial parameters in a single trial. Black dot represents the projections of initial fixation/gaze and the blue dot corresponds to final gaze (G). The red dot depicts the location of target (T) in relation to the original landmark (L) position (broken intersecting lines), and the green dot represents the virtually shifted target (T’) fixed to the shifted landmark (L’) location (solid intersecting lines). Note that the final gaze was placed between the T and T’ (i.e., it shifted toward the shifted landmark). In head-unrestrained conditions, the target can be encoded in egocentric coordinates (eye, head or body) (Sajad et al., 2015). We plotted two continuums: a T-G continuum (egocentric, we divided the space between T and G into ten equal steps, thus treating these spatial codes as a continuous spatiotemporal variable) to compute the gaze influence and an allocentric shift T-T’ continuum (original target to virtual target) based on the landmark shift to compute the influence of landmark shift on the neuronal activity. **(B)** A logical schematic of response field analysis. X-axis depicts the coordinate frame, and the Y-axis corresponds to the related activity to the target. Simply, if the activity related to a fixed target is plotted in the correct reference frame, this will lead to the lowest residuals, i.e., if the neural activity to a fixed target location is fixed (left), then the data (blue) would fit (pink) better on that, leading to lower residuals in comparison with when the activity for the target is plotted in an incorrect frame, thus leading to higher residuals (right).

Further, to quantify the influence of the landmark shift, we created **(Fig. 4A**, lower right) another continuum (T-T’, the line between the original target and the target if it were fixed to the landmark) between Te (Target fixed in eye coordinates) and T’e (virtual Target fixed in landmark coordinates), computed in eye coordinates with ten intermediary steps between, and additional steps on the flanks (10 beyond Te and 10 beyond T’e). These additional ten steps beyond the canonical models were included 1) to quantify if neurons can carry abstract spatial codes around the canonical models and 2) to eliminate the misleading edge effects (else the best spatial models would cluster around the canonical models). The T-T’ continuum will allow us to test whether SEF code is purely egocentric (based on T), allocentric (based on T’), or contains the integrated allocentric + egocentric information that eventually actuates the behavior. Note that AW and T-T’ are geometrically similar, but the first describes behavioral data whereas the second describes neural data. Hereafter, we will refer to T-T’ as the ‘allocentric shift’.

### Fitting Neural Response Fields against Spatial Models

In order to test different spatial models, they should be spatially separable (Keith et al. 2009; Sajad et al. 2015). The natural variation in monkeys’ behavior allowed this spatial separation (see Results for details). For instance, the variability induced by memory-guided gaze shifts permitted us to discriminate target coding from the gaze coding; the initial eye and head locations allowed us to distinguish between different egocentric reference frames and variable eye and head movements for a gaze shift permitted the separation of different effectors. As opposed to the decoding methods that generally test if a spatial characteristic is implicitly coded in patterns of population activity (Brandman et al., 2017; Bremmer et al., 2016), the method employed in this study method directly examines which model best predicts the activity in spatially tuned neural responses. The logic of our response field fitting in different reference frames is schematized in **Fig. 4B**. Precisely, if the response field data are plotted in the correct best/reference frame, this will yield the lowest residuals (errors between the fit and data points) compared with other models, i.e., if a fit computed to its response field matches the data, then this will yield low residuals **(Fig. 4B**,left). On the other hand, if the fit does not match the data well, this will lead to higher residuals **(Fig. 4B**, right). For example, if an eye-fixed response field is computed in eye-coordinates this will lead to lower residuals and if it is plotted in any other inferior/incorrect coordinate, such as space coordinate, this will produce higher residuals (Sajad et al., 2015).

In reality, we employed a non-parametric fitting method to characterize the neural activity with reference to a spatial location and also varied the spatial bandwidth of the fit to plot any response field size, shape, or contour (Keith et al., 2009). We tested between various spatial models using Predicted Residual Error Some of Squares (PRESS) statistics. To independently compute the residual for a trial, the actual activity related to it was subtracted from the corresponding point on the fit computed over all the other trials (like cross-validation). Notably, if the spatial physical shift between two models results in a systematic shift (direction and amount), this will appear as a shifted response field or expanded response field and our model fitting approach would not be able to distinguish two models as they would produce virtually indistinguishable residuals. Because in our study, the distribution of relative positions across different models also possesses a non-systematic variable component (e.g., variability in gaze endpoint errors, or unpredictable landmark shifts), the response fields invariably stayed at the same location, but the dissociation between spatial models was based on the residual analysis.

We plotted response fields (visual and movement) of neurons in the coordinates of all the canonical (and intermediate) models. To map the visual response in egocentric coordinates, we took eye and head orientations at the time of target presentation, and for movement response fields we used behavioral measurements at the time when the gaze shift started (Keith et al. 2009; DeSouza et al. 2011; Sajad et al. 2015). Likewise, for the allocentric models, we used the initial and the shifted landmark location to plot our data. Since we did not know the size and shape of a response field a priori and since the spatial distribution of data was different for every spatial model (e.g. the models would have a smaller range for head than the eye models), we computed the non-parametric fits with different kernel bandwidth (2-25°), thus making sure that we did not bias the fits toward a particular size and spatial distribution. For all the tested models with different non-parametric fits, we computed the PRESS residuals to reveal the best model for the neural activity (that yielded the least PRESS residuals). We then statistically (Keith et al., 2009) compared the mean PRESS residuals of the best model with the mean PRESS residuals of other models at the same kernel bandwidth (two-tailed Brown-Forsythe test). Finally, we performed the same statistical analysis (Brown-Forsythe) at the population level (Keith et al., 2009; DeSouza et al., 2011). For the models in intermediate continua, a similar procedure was used to compute the best fits.

### Testing for Spatial Tuning

The above described method assumes that neuronal activity is structured as spatially tuned response fields. This does not imply that other neurons do not sub-serve the overall population code (Bharmauria et al., 2016; Chaplin et al., 2018; Goris et al., 2014; Leavitt et al., 2017; Pruszynski and Zylberberg, 2019; Zylberberg, 2018) but with our method only tuned neurons can be explicitly tested. We tested for the neuronal spatial tuning as follows. We randomly (100 times to obtain random 100 response fields) shuffled the firing rate data points across the position data that we obtained from the best model. The mean PRESS residual distribution (PRESS_random_) of the 100 randomly generated response fields was then statistically compared with the mean PRESS residual (PRESS_best-fit_) distribution of the best-fit model (unshuffled, original data). If the best-fit mean PRESS fell outside of the 95% confidence interval of the distribution of the shuffled mean PRESS, then the neuron’s activity was deemed spatially selective. At the population level, some neurons displayed spatial tuning at certain time-steps and others did not because of low signal/noise ratio. Thus, we removed the time steps where the populational mean spatial coherence (goodness of fit) was statistically indiscriminable from the baseline (before target onset) because there was no task-related information at this time and thus neural activity exhibited no spatial tuning. We defined an index (Coherence Index, CI) for spatial tuning. CI for a single neuron, which was calculated as (Sajad et al., 2016):

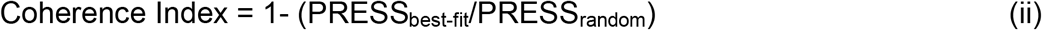

If the PRESS_best-fit_ was similar to PRESS_random_ then the CI would be roughly 0, whereas if the best-fit model is a perfect fit (i.e., PRESS_best-fit_ = 0), then the CI would be 1. We only included those neurons in our analysis that showed significant spatial tuning.

### Time-normalization for spatiotemporal analysis

A major aim of this study was to track the progression of the T-G and T-T’ codes in spatially tuned neuron populations, from the landmark-shift until the saccade onset. The challenge here was that this period was variable, so that averaging data across trials for each neuron would result in the mixing of very different signals (see extended **Figure 11-1)**. To compensate for this, we time-normalized this data (Sajad et al. 2016; Bharmauria et al. 2020). To do this, the neural firing rate (in spikes/second; the number of spikes divided by the sampling interval for each trial) was sampled at 7 half-overlapping time windows (with a range of 80.5 – 180.5 ms depending on trial duration). The rationale behind the bin number choice was to make sure that the sampling time window was wide enough, and therefore robust enough to account for the stochastic nature of spiking activity (thus ensuring that there were sufficient neuronal spikes in the sampling window to do effective spatial analysis) (Sajad et al., 2016). Once the firing rate was estimated for each trial at a given time-step, they were pooled together for the spatial modeling. Note that the final (7th) time step also contained some part of the perisaccadic sampling window. Finally, we performed our T-G / T – T’ fits on the data for each of these time bins. In short, this procedure allowed us to treat the whole sequence of memory-motor responses from the landmark shift until saccade onset as a continuum.

### Statistical Table 1

**Statistical Table 1:**
| ANALYSIS | STATISTICAL TEST | POWER |
| --- | --- | --- |
| Distribution significantly shifted from a point | One-sampled Wilcoxon signed ranked test | $p < 0.05$ |
| Comparison between models | Brown-Forsythe test | $p = 10^0 = 1$; the best model |
| Comparison between distributions | Mann-Whitney U-test | $p < 0.05$ |
| Transformation between the visual and the motor responses for visuomotor neurons | Wilcoxon matched-pairs signed rank | $p < 0.05$ |
| Spatial tuning test for a neuron | Coherence index (see above) | $p < 0.05$ |
| Comparison between visuomotor and motor neurons in SEF and FEF | 3-factor ANOVA analysis | $p < 0.05$ |
| Correlation between T-G and T-T' | Spearman Correlation | $p < 0.05$ |

## RESULTS

### Influence of Landmark shift on behavior

To investigate how the landmark influences behavior toward an object of interest, we used a cue-conflict task, where a visual landmark shifted during the memory delay, i.e., between the target presentation and the gaze shift **(Fig. 1A)**. The task is further schematized in **Figure 1B** with possible target, original landmark, virtual target and shifted landmark locations. We then computed the influence of the landmark shift as described previously (Li et al., 2017) and further confirmed it in the current dataset. This is computed as allocentric weight (AW), i.e., the component of gaze end points along the axis between the target and the cue shift direction. If AW equals 0, then it implies no influence of the shifted landmark, whereas if AW equals 1 then it indicates a complete influence of landmark shift on the gaze (see methods). Note that we performed this analysis on the same trials as used for the analysis of spatially tuned activity described below.

In both animals (Monkey V and Monkey L), the gaze endpoints scattered along the one-dimensional axis of the landmark shift **(Fig 1C)**, producing a bell-like distribution of gaze errors. However, both distributions showed a highly significant (p < 0.001, one sampled Wilcoxon signed rank test) shift from the original target (0) in the direction of the landmark shift (with a mean AW of 0.36 in Monkey V and 0.29 in Monkey L). There was considerable trial-to-trial variance around these means, but note that such variance was present or even larger in the absence of a landmark (Li et al., 2017; Sajad et al., 2015). The overall average error (distance) between T and final gaze, i.e., including errors in all directions, was 8.19 ± 5.27 (mean ± SD) and 10.35 ± 5.89 for Monkey V and Monkey L, respectively. No correlation was found between the AW as a function of saccade latency (Bharmauria et al., 2020). These results generally agree with previous cue-conflict landmark studies (Byrne and Crawford, 2010; Fiehler et al., 2014; Neggers et al., 2005) and specifically with our use of this paradigm in the same animals, with only slight differences owing to collection of data on different days (Bharmauria et al., 2020; Li et al., 2017).

In order to determine when the landmark first influenced the SEF spatial code, and when that influence became fully integrated with the default SEF codes, we asked the following questions in their logical order: 1) What are the fundamental egocentric and/or allocentric codes in the SEF 2) How does the landmark shift influence these neural codes 3) What is the contribution of different SEF cell types (visual, visuomotor, and motor), and finally, 4) is there any correlation between the SEF allocentric and egocentric codes, as we found in the FEF?

### Neural recordings: general observation and classification of neurons

To understand the neural underpinnings of the behavior (as revealed above) in the SEF activity, we recorded visual and motor responses from over 200 SEF sites using tungstenmicroelectrodes, while the monkey performed the cue-conflict task **(Fig 2A-B)**. We analyzed a total of 256 neurons and after applying our rigorous exclusion criteria (see methods), we were finally left with 68 significantly spatially tuned neurons (see methods): 32 significantly spatially tuned visual responses (including V and VM) and 53 (including M and VM) significantly spatially tuned motor responses. Many other neurons (n = 188) were not spatially tuned or did not respond to the event and were thus excluded from further analyses. Typically, neurons that were not spatially tuned were responsive at some time during the task throughout both visual fields, and did not show preference for any of the spatial models that we described below.

The mean spike density graphs (with the 100-ms sampling window) for the visual (red) and motor (black) neurons are shown in **Figure 2C** and **2D** respectively. We then performed the non-parametric analysis (see **Fig. 4A-B**) while using all trials throughout the response field of each neuron (dark black and red curves), but we also display the top 10 % from every neuron during the sampling window (light red and gray for visual and motor activity respectively), roughly corresponding to the ‘hot-spot’ of the response field. We first employed our model-fitting approach to investigate SEF spatial codes, starting (since this has not been done with SEF before) with a test of the most fundamental models.

### Visual activity fitting in egocentric and allocentric spatial models

We began by testing all the egocentric and allocentric models **(Fig. 3)** in the sampling window (80-180 ms relative to target onset) of the visual response (including pure visual and visuomotor neurons). **Figure 5** shows a typical example of analysis of a visual response field. **Figure 5A** displays the raster and spike density (pink curve) plot of a visually responsive neuron aligned to the target onset (blue line as indicated by downward blue arrow). The shaded pink area corresponds to the sampling window for response field analysis. **Figure 5B** shows the closed (spatially restricted) response field for this neuron in the original target-in-space coordinates (Ts) corresponding to actual stimulus locations. Each circle corresponds to the magnitude of response in a single trial. The heat map represents the non-paramedic fit to the neural data, with residuals (difference between data and fit) plotted to the right. This neuron has a hot-spot (red) near the center of the response field.

**Figure 5.**
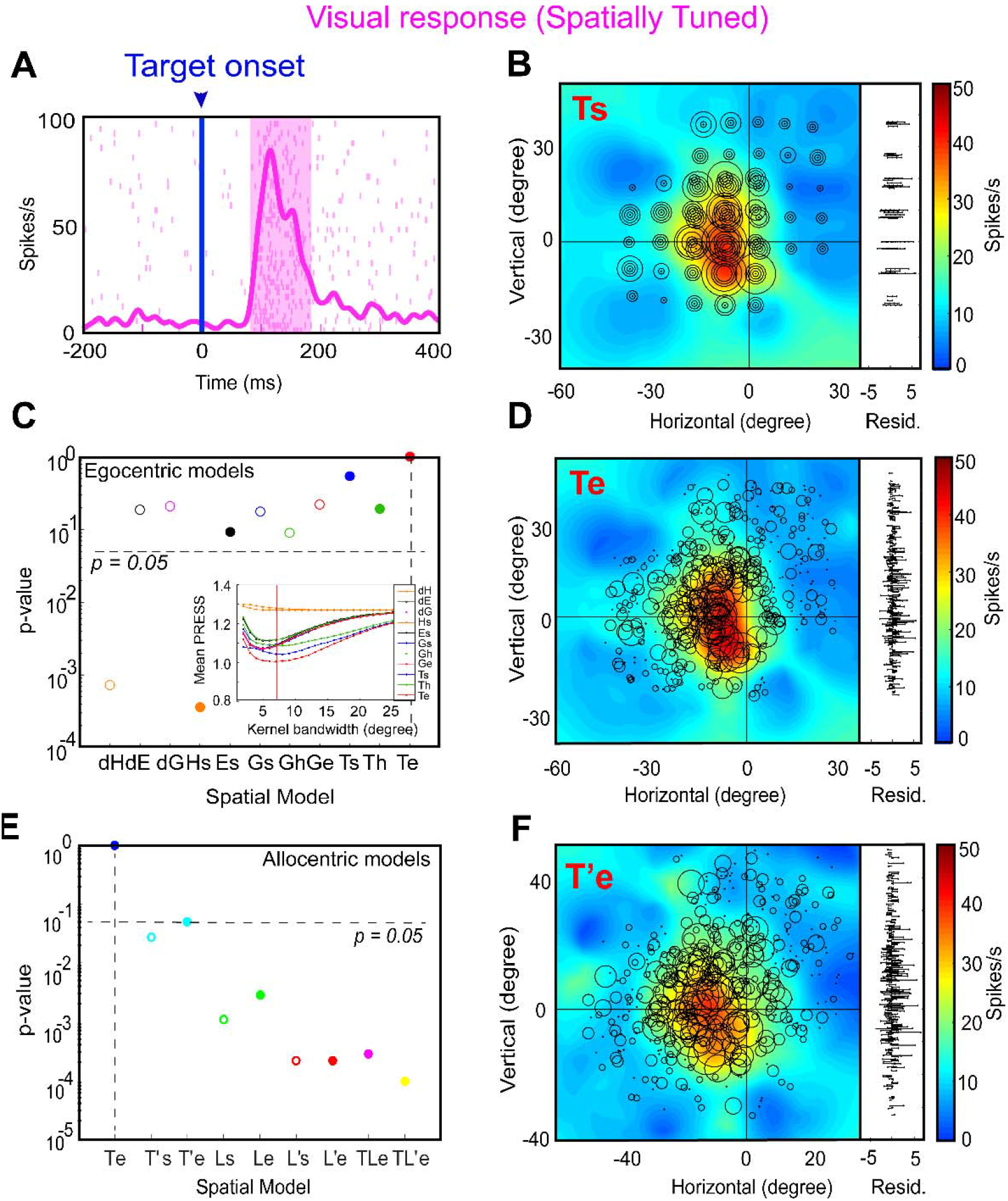
An example of response field analysis of a visual neuron. **(A)** Raster/spike density plot (with top 10 % responses) of the visual neuron aligned to the onset of target (blue arrow); the shaded pink region corresponds to the sampling window (80-180 ms) for response field analysis. **(B)** Representation of neural activity for Ts: target in space (screen). The circle corresponds to the magnitude of the response and heat map represents the non-parametric fit to these data (red blob depicts the hot-spot of the neuronal response field). The corresponding residuals are displayed to the right. To independently compute the residual for a trial, its activity was subtracted from the point corresponding to the fit computed on the rest of the trials. The residuals are one-dimensional for every data point from the fit projected onto the vertical axis for illustrative purposes. **(C)** The p-values statistics and comparison between Te (p = 10^0^ = 1; the best-fit spatial model that yielded lowest residuals) and other tested models (Brown-Forsythe test). The inset shows the mean residuals from the PRESS-statistics for all spatial models at different kernel bandwidths (2-25°). The red vertical line depicts the lowest mean PRESS for Te (target in eye coordinates) with a kernel bandwidth of 7. **(D)** Representation of the neural activity in Te (target relative to eye), the best coordinate. **(E)** The p-value statistics and the comparison of the best egocentric model, Te, with other allocentric spatial models. Te is still the best-fit model. **(F)** Representation of the neural activity in T’e, i.e., shifted target relative eye. 0, 0 represents the center of the coordinate system that led to the lowest residuals (best fit).

**Figure 5C** provides a statistical summary (p-values) comparing the goodness of fits for each of our canonical egocentric models (x-axis, from **Figure 3A**) relative to the model with the lowest residuals. For this neuron, the response field was best fit across target in eye coordinates (Te) with a Gaussian kernel bandwidth of 6° (see inset, the red vertical line indicates the best kernel bandwidth). Thus, Te is the preferred egocentric model for this neuron, although most other models were not significantly eliminated. **Figure 5D** shows the response field plot of the neuron in Te coordinates. Comparing Ts and Te, one can see how larger circles (larger bursts of activity) tend to cluster more together at the response field hot spot in Te than Ts, and the residuals look (and were) smaller, implying that Te is the best coordinate system for this neuron. The bottom row of **Figure 5** shows a similar analysis of the same neuron with respect to our allocentric models (from **Fig. 3B**). **Figure 5E** provides a statistical comparison (p-values) of the residuals of these models relative to Te, which we kept as a reference from the egocentric analysis. Te still provided the lowest residuals, whereas nearly every allocentric model was statistically eliminated. The exception was T’e (target-in-eye coordinates but shifted with the landmark) perhaps because it was most similar to Te. We have used this coordinate system for the example response field plot in **Figure 5F**. An example of a spatially untuned neuron is shown in **Figure 5-1**.

**Figure 6** shows the pooled fits for visual (Visual + Visuomotor) responses across all spatially tuned neurons (n = 32) for both egocentric (left column) and allocentric (right column) analyses. In each column, the top row provides the distributions of mean residuals, the middle the p-value comparisons, and the bottom row the fraction of spatially tuned neurons that preferred each model. In the egocentric analysis **(Fig. 6A-B),** Te was still the best fit overall (and was preferred in ~40% of the neurons). The head-centered models and most of the effector-specific models were statistically eliminated but dE, dG, Ge and Ts were not eliminated. Compared with the allocentric models **(Fig. 6C-D),** Te performed even better, statistically eliminating all other models. Overall, these analyses of egocentric models suggest that Te was the predominant code in the SEF visual responses, just as we found for the FEF in the same task (Bharmauria et al., 2020) and both the FEF and SC in a purely egocentric gaze task (Sadeh et al., 2015; Sajad et al., 2015).

**Figure 6.**
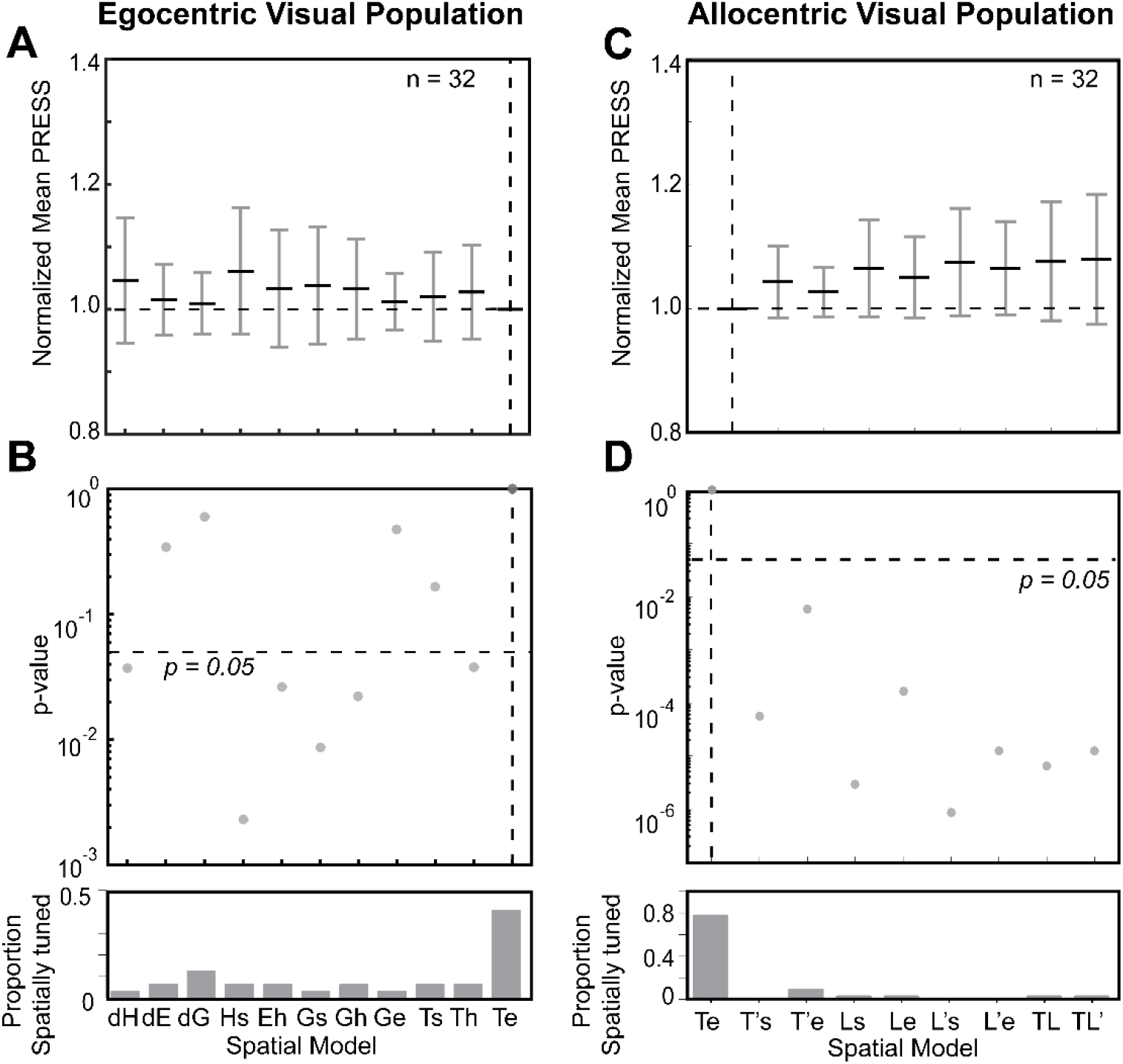
Egocentric and allocentric population fit residuals and statistics for all visual neurons. **(A)** The mean mean PRESS (± SEM) residuals of all visually responsive neurons (n = 32), i.e., data relative to fits computed to all tested egocentric models. These values were normalized by dividing by the mean PRESS residuals of the best spatial model, in this case Te. **(B)** P-Value statistics performed on the residuals shown above (Brown-Forsythe test). Te (broken vertical blue line) is the best fit, however dE, dG, Ge and Ts were also retained. (C) Same as **A** but for best-fit egocentric model comparison with all allocentric models. (D) Same as **B**, Te is still the best fit.

### Motor activity fitting in egocentric and allocentric spatial models

We then proceeded with the analysis of neural activity for motor responses (including the motor and visuomotor neurons) i.e., neurons firing in relation to the saccade onset. Despite the landmark-induced shifts and variable errors in gaze as described above, overall there was still a strong correlation between target direction and final gaze position (Bharmauria et al., 2020), and thus motor response fields tended to align with sensory response fields. **Figure 7** shows a typical analysis of a motor neuron. **Figure 7A** displays the raster and spike density plot for a motor response (the shaded area corresponds to the sampling window and **Figure 7B** shows the corresponding response field, plotted in Ts coordinates. **Figure 7C** provides a statistical analysis of this neuron’s activity against all egocentric spatial models (same convention as **Figure 5**). Here, Ge (future gaze direction relative to initial eye orientation) was the preferred model for this neuron, although several other models were also retained. **Figure 7D** shows the corresponding data in Ge coordinates, where again, residuals are lower and similar firing rates cluster. **Figure 7E** shows the statistical testing of different allocentric models using Ge as the egocentric reference. Once again, the egocentric model ‘wins’, eliminating all of the allocentric models except for T’e (used in the example response field plot, **Fig. 7F**).

**Figure 7.**
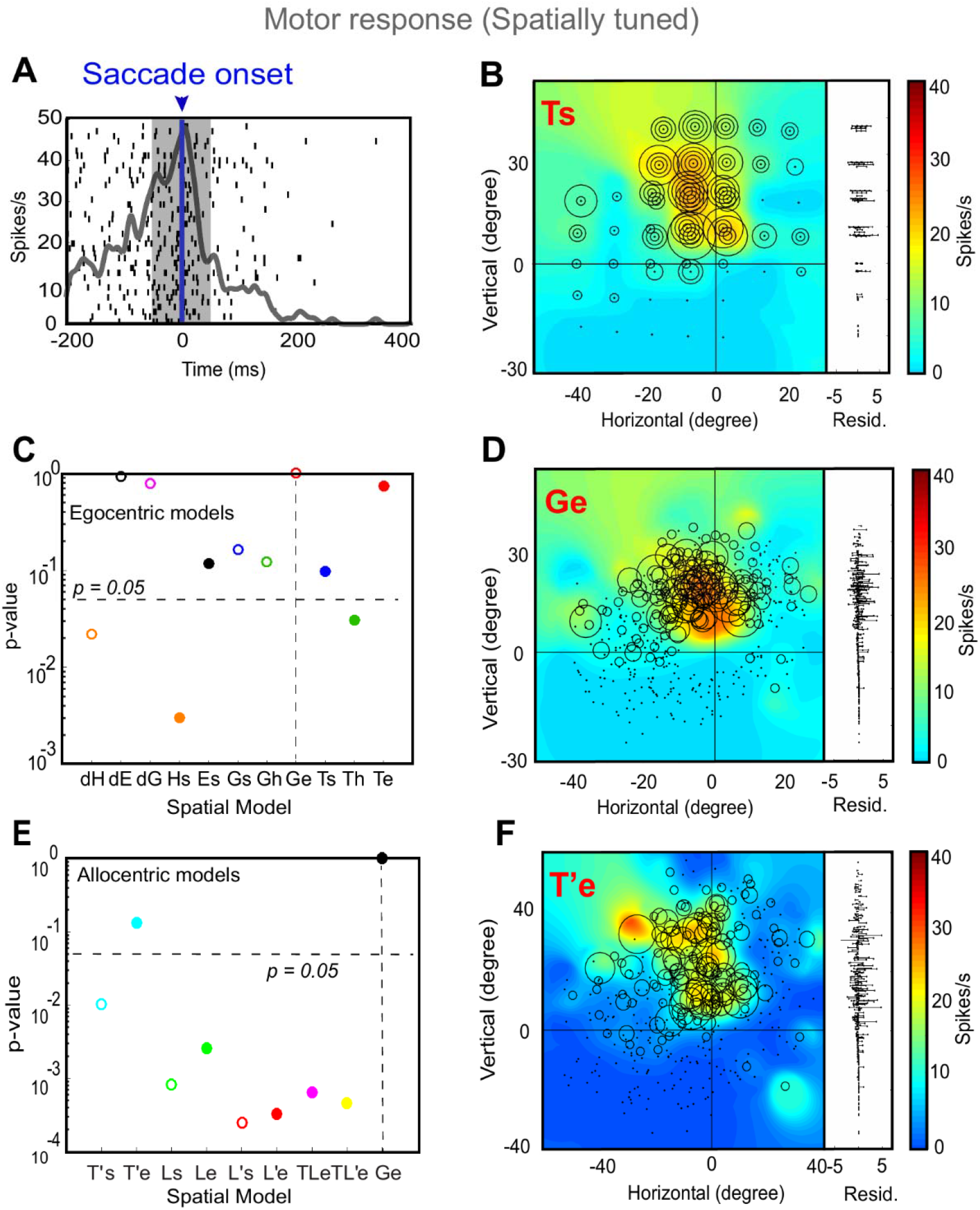
An example of response field analysis of a motor response. **(A)** Raster/spike density plot (with top 10 % responses) of the motor neuron aligned to the saccade onset (blue arrow); the shaded grey area corresponds to the sampling window (−50-50 ms) for response field analysis. **(B)** Representation of neural activity for Ts: target in space (screen). The corresponding residuals are displayed to the right. **(C)** The p-values statistics and comparison between Ge (p = 10^0^ = 1; the best-fit spatial model that yielded lowest residuals) and other tested models (Brown-Forsythe test). **(D)** Representation of the neural activity in Ge (future gaze relative to eye), the best coordinate. **(E)** The p-value statistics and the comparison of the best egocentric model, Ge, with other allocentric spatial models. Ge is still the best-fit model, with T’e (shifted target in eye coordinates) as the second best model implying the influence of the shifted landmark on the motor response. **(F)** Representation of the neural activity in T’e, i.e., shifted target relative eye. 0, 0 represents the center of the coordinate system that led to the lowest residuals (best fit).

**Figure 8** summarizes our population analysis of motor responses (n = 53, Motor + Visuomotor) against our egocentric and allocentric models, using the same conventions as **Figure 6**. Overall, of the egocentric models, dE yielded the lowest residuals, but dE was statistically indistinguishable from dG and Ge, which also yielded similar amounts of spatial tuning **(Fig. 8A**). Importantly, Te was now eliminated, along with dH, Hs, Ts, and Th **(Fig. 8B).** We retained Ge as our egocentric reference for comparison with the allocentric fits, because it is mathematically similar to Te (used for the visual analysis), and has usually outperformed most motor models in previous studies (Bharmauria et al., 2020; Klier et al., 2001; Martinez-Trujillo et al., 2004; Sadeh et al., 2015; Sajad et al., 2015). This time, comparisons with Ge statistically eliminated all of our allocentric models at the population level. In summary, these analyses suggest that Ge (or something similar), and not Te, was the preferred model for motor responses, as reported in our studies on FEF and SC (Bharmauria et al., 2020; Sadeh et al., 2015; Sajad et al., 2015).

**Figure 8.**
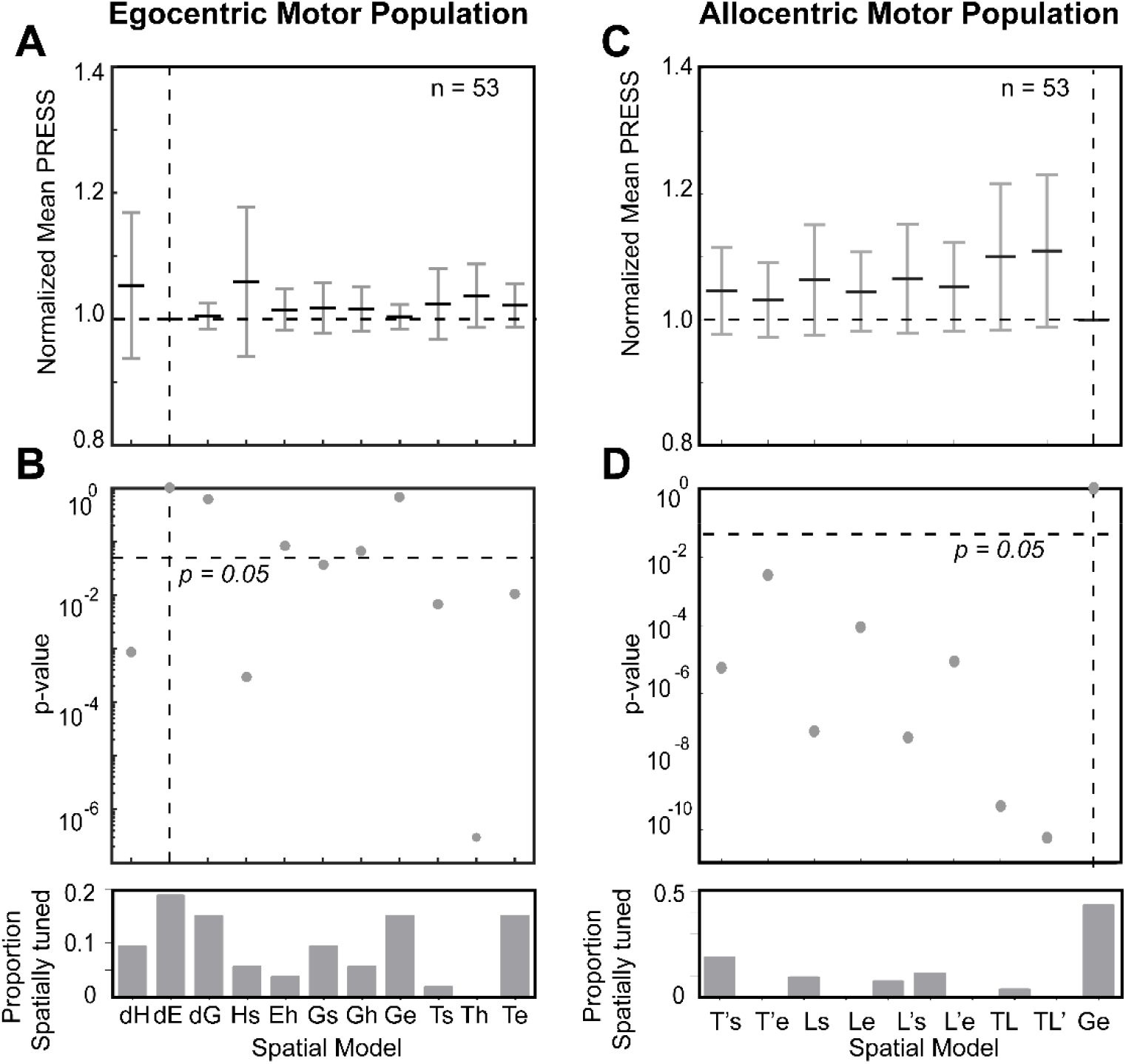
Egocentric and allocentric population fit residuals and statistics for all motor neurons. **(A)** The mean mean PRESS (± SEM) residuals of all motor responses (n = 53), i.e., data relative to fits computed to all tested egocentric models. These values were normalized by dividing by the mean PRESS residuals of the best spatial model, in this case dE. **(B)** P-Value statistics performed on the residuals shown above (Brown-Forsythe test). dE (broken vertical blue line) is the best fit, however dG and Ge were also retained and very close to dE. **(C)** Same as **A** but for best-fit egocentric model comparison with all allocentric models. **(D)** Same as **B**, Ge is the best fit.

### Visuomotor transformation along the intermediate frames: T-G and T-T’ continua

Thus far, we found that SEF continued to be dominated by eye-centered target (T) and gaze (G) codes, like other saccade-related areas (FEF and SC). However, it is possible that the actual codes fall within some intermediate code, as we have found previously in the FEF (Bharmauria et al., 2020). Therefore, as described in the Methods section **(Fig. 4)** and **Figure 9**, we constructed the same spatial continua to quantify the detailed sensorimotor transformations in the SEF: a T-G (specifically Te-Ge) continuum to quantify the amount of transition from target to future gaze coding, and a T-T’ continuum (specifically Te-T’e; similar to our behavioral AW score) to quantify the influence of the landmark on the target code. Following analysis shows an example of a visual and a motor response.

**Figure 9.**
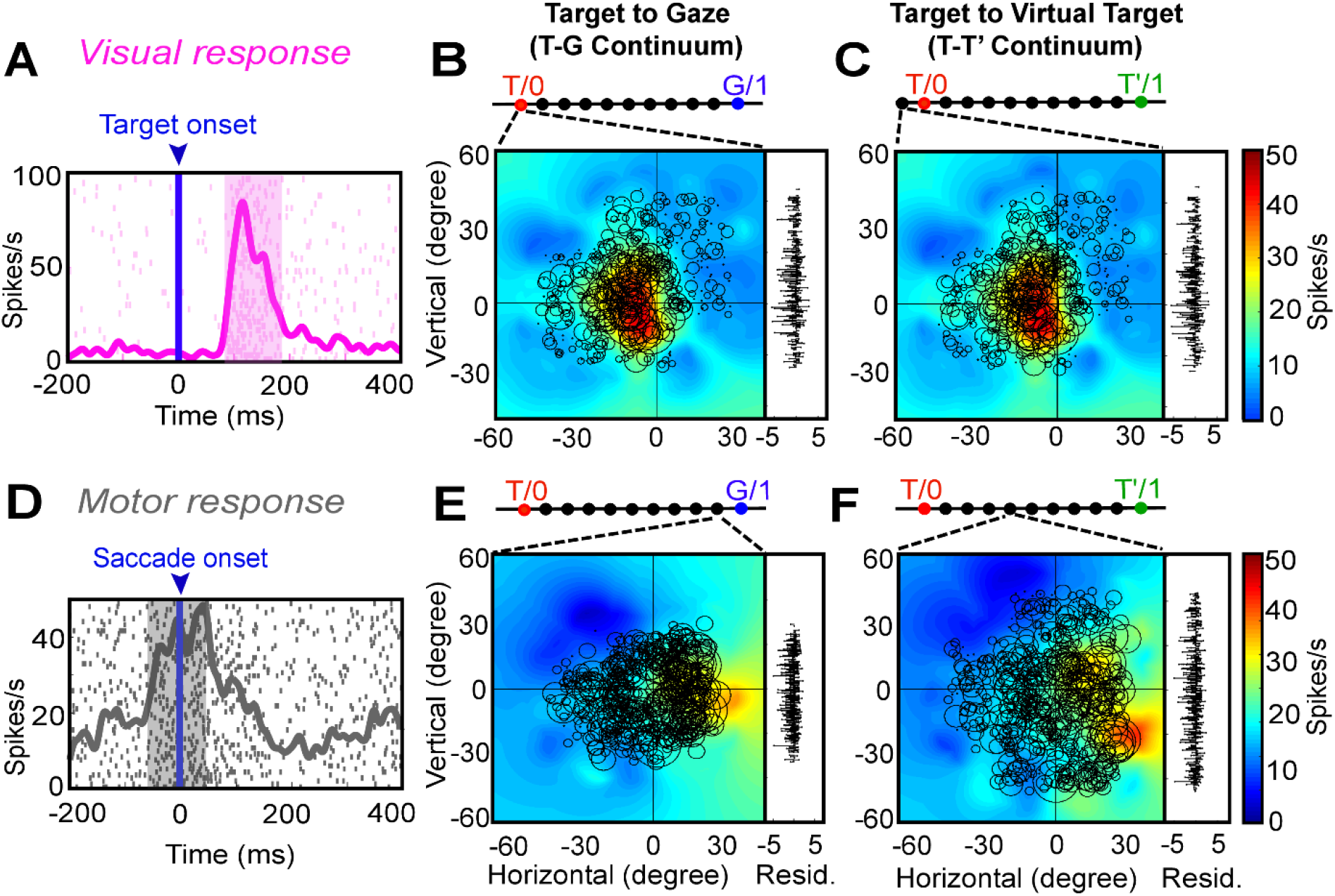
Egocentric (target to gaze, T-G continuum) and allocentric (target to virtually shifted target, T-T’ continuum) **(A)** Raster/spike density plot (with top 10 % responses) of the visual neuron (same neuron as **Fig 6**) aligned to the target onset (blue arrow). **(B)** Representation of the response field in the eye-centered coordinates obtained from the best-fit of the neural data along the T-G continuum (bar above the response field). To the right of the plot are the residuals between the individual trial data and the response field fit. The response field was best located exactly at T **(C)** Response field along the T-T’ continuum, the best fit is located one step beyond T indication no influence of the landmark shift. **(D)** Raster/spike density plot of a motor response aligned to the onset of the saccade (blue arrow) **(E)** Representation of the motor response field along the T-G continuum, the response field fits best at the ninth step (converging broken lines) from T (one step to G). **(F)** Representation of the motor response field along the T-T’ continuum, the response field fits best at 4^th^ step from T, indicating the influence of the landmark shift on the motor response.

**Figure 9A** shows the raster and the spike density plot for a visually responsive neuron aligned to the target onset (same neuron as **Figure 5**). **Figure 9B** displays the best-fit response field plot of the neuron along the T-G continuum, where the circle represents the magnitude of the response, the heat map represents the non-parametric fit to the data, and the residuals are plotted to the right. The converging broken lines pointing to bar at the top represent the corresponding point of best fit along the ten equal steps between T and G. Here, the response field of neuron fits best exactly at T as indicated by the broken lines. **Figure 9C** shows the response field plot of the same data along the T-T’ continuum. Here, the best fit for the response field was located only one step (10% beyond T, in the direction away from T’) demonstrating no influence of the future landmark shift on the initial visual response.

What then happens after the landmark shift? **Figure 9D** depicts the raster and spike density plot of a motor neuron, aligned to the saccade onset. Along the T-G continuum **(Figure 9E),** the best response field fit was at the 9^th^ step, i.e., 90% toward G (suggesting a near-complete transformation to gaze coordinates), whereas the best fit along the T-T’ continuum **(Figure 9F),** was at 4^th^ step from T, i.e., 40% toward T’ (suggesting landmark influence similar to that seen in our behavioral measure).

### Population analysis along the T-G and T-T’ continuum

How representative were the examples shown above of our entire population of data? To answer that question, we performed the same analysis on our entire population of spatially tuned visual (both visual and visuomotor) and motor (both visuomotor and motor) responses. **Figure 10** shows the distribution of best fits for visual responses (top row) and motor responses (bottom row) neurons. T-G distribution of visually responding neurons **(Figure 10A)** showed a primary cluster and peak around T, but overall was shifted slightly toward G (mean = 0.3; median = 0.15), because of a smaller secondary peak at G. This suggests that most visual responses encoded the target, but some already predicated the future gaze location. This is similar to what has been reported in FEF too (Bharmauria et al., 2020; Sajad et al., 2015). The motor distribution (**Figure 10B**, n = 53) showed the opposite trend: a smaller cluster remained at T, but the major cluster and peaks were near G. Overall, this resulted in a significant (p < 0.0001, one sampled Wilcoxon signed rank test) shift toward G (mean = 0.72; median= 0.8). Notably, the motor and visual distributions were significantly different from each other (p < 0.0001; Mann-Whitney U-test).

**Figure 10.**
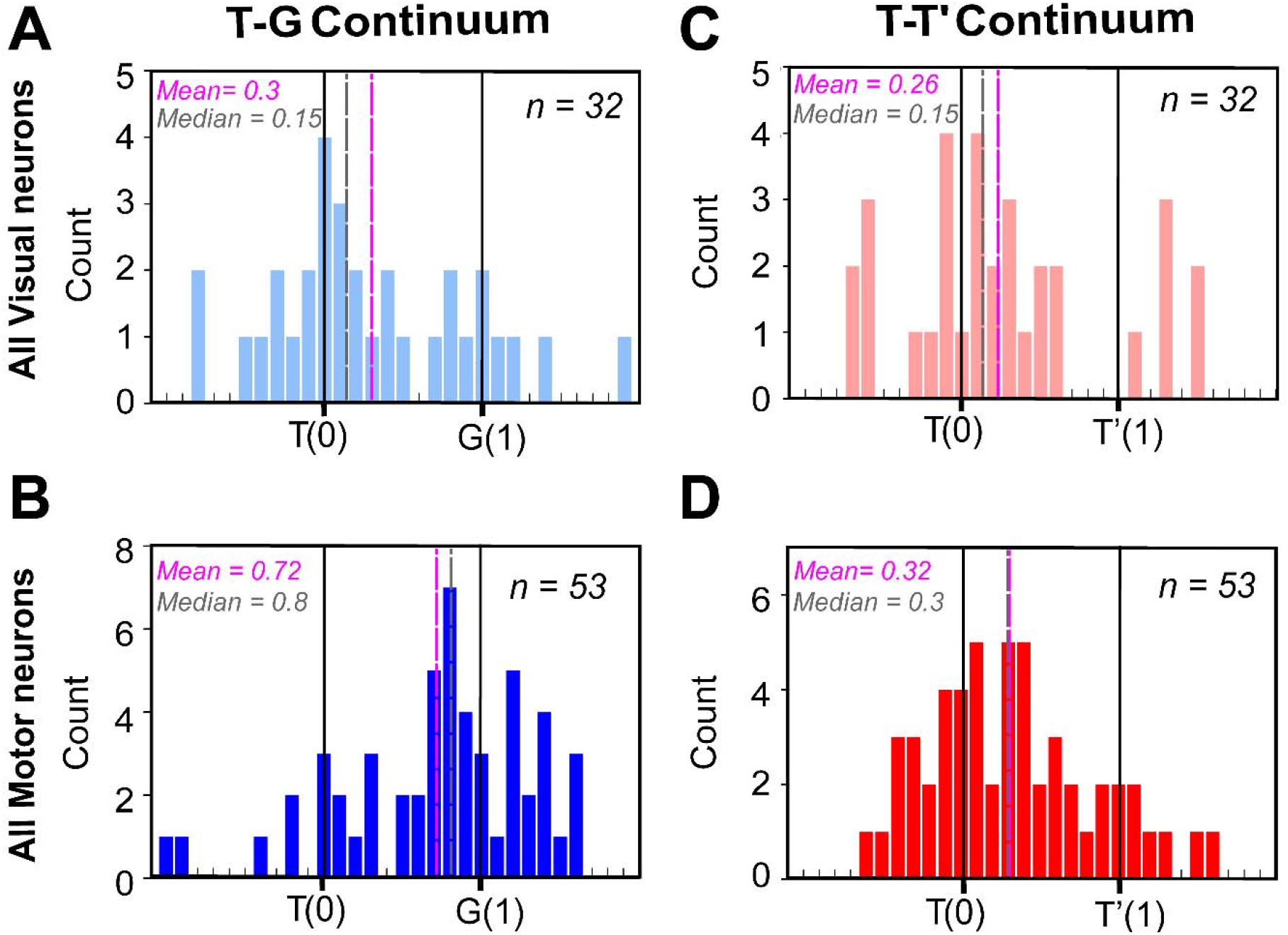
Frequency distribution along the T-G and T-T’ continua at the population level. (**A)** Frequency distribution of all spatially tuned visual responses (n = 32) along the T-G continuum with the best fit closer to T (mean = 0.3; median = 0.15). **(B)** Frequency distribution of all spatially tuned motor responses (n = 53) along the T-G continuum with a significantly shifted distribution toward G (mean = 0.72; median = 0.8; p < 0.0001, one sampled Wilcoxon signed rank test). **(C)** Frequency distribution of all spatially tuned visual responses (n = 32) along the T-T’ continuum with the best fit closer to T (mean = 0.26; median = 0.15). **(D)** Frequency distribution of all spatially tuned motor responses (n = 53) along the T-T’ continuum with a significantly shifted distribution toward T’ (mean = 0.32; median = 0.3; p = 0.0002, one sampled Wilcoxon signed rank test) indicating the influence of the landmark shift on the motor responses.

Along the T-T’ continuum **(Fig. 10C),** the best fits for the visual population peaked mainly around T, but overall showed a small (mean = 0.26; median = 0.15) but non-significant shift toward T’ (p = 0.07, one sampled Wilcoxon signed rank test). The motor population **(Fig. 10D)** shifted further toward T’ (mean = 0.32; median= 0.3). This overall motor shift was not significantly different from the overall visual population (p = 0.53; Mann-Whitney U-test), but it was significantly shifted from T (p = 0.0002, one sampled Wilcoxon signed rank test). In general, this T-T’ shift resembled the landmark influence on actual gaze behavior. Notably, at the single cell level there was a significant T-G transition **(Figure 10-1)** between the visual and the motor responses *within* visuomotor neurons (n = 16; p = 0.04, Wilcoxon matched-pairs signed rank test), but not along the TT’ continuum (n = 16; p = 0.32, Wilcoxon matched-pairs signed rank test).

Overall, this demonstrates a target-to-gaze transformation similar to the SC and FEF (Sadeh et al., 2015; Sajad et al., 2015) and a similar significant landmark influence in the motor response as we found in FEF (Bharmauria et al., 2020). However, we do not yet know how different cell types contribute to this shift and if this landmark influence has some relationship to the egocentric (T-G) transformation as revealed in FEF (Bharmauria et al., 2020).

### Contribution of different cell types and the allocentric shift

We next examined how different neuronal classes are implicated in the landmark influence as noticed above **(Fig. 11).** To this goal, as in our FEF study on the same task (Bharmauria et al., 2020), we focused our analysis on a 7-step time-normalized analysis aligned to onset of the landmark-shift until the saccade onset **(Figure 11-1).** Since the second delay was variable, the time-normalization procedure allowed us to treat the corresponding neural activity as a single temporal continuum (Bharmauria et al., 2020; Sajad et al., 2016). By employing this procedure on the neural activity, we tracked the progression of T-G and T-T’ continua to quantify the gaze and landmark-shift influence, respectively.

**Figure 11.**
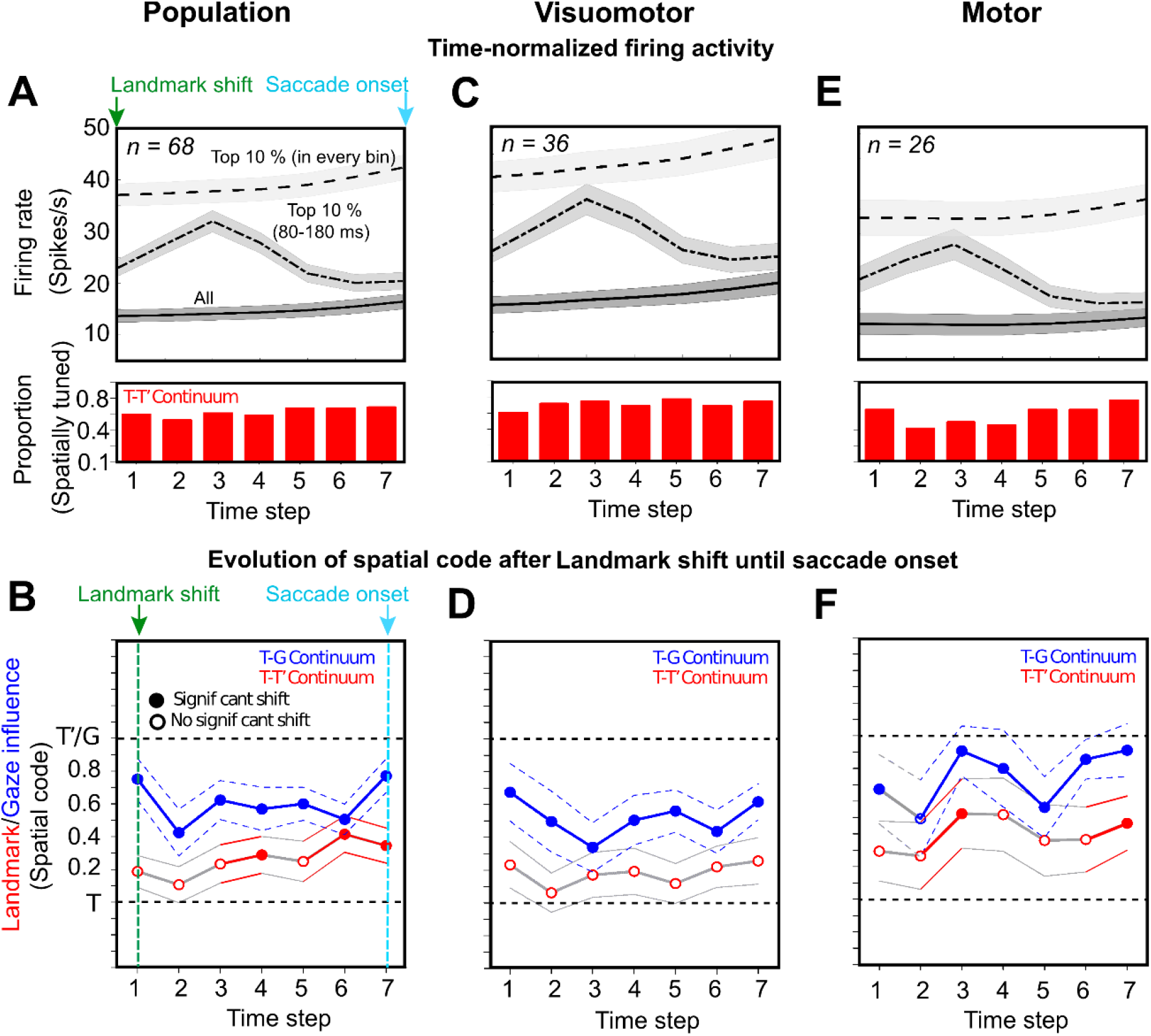
Spatiotemporal analysis aligned to landmark shift until saccade onset. **(A, B)** Spatiotemporal analysis of whole of the population (n = 68) aligned to the landmark onset (green arrow) unit the saccade onset (cyan arrow). **(A)** Time-normalized neural activity divided into 7 half-overlapping bins for all trials (bottom trace), top 10 % trials in each bin (top trace), and top 10 % trials in 80-180 ms window aligned to landmark-shift (middle trace). **(B)** Progression of the spatial code for all neurons (n = 68) along the T-G and T-T’ continua. **(C)** Same as **A** but for visuomotor neurons (n = 36). **(D)** Progression of spatial code for visuomotor neurons along the T-G and T-T’ continua. None of the steps displayed a significant shift toward T’ along the T-T’ continuum. **(E)** Same as **A** but for motor neurons (n = 26). **(F)** Progression of the spatial code in time for motor neurons along the T-G and T’-T’ continua. A significant shift toward T’ was noted along the T-T’ continuum at the 3rd step and the 7^th^ step (just before the saccade onset). Note: the histogram below the spike density plots displays the proportion of spatially tuned neurons at each time-step.

**Figure 11A** displays the mean activity of the entire spatially tuned population (n = 68) of neurons divided into 7 time-normalized bins from landmark-shift onset to the saccade onset (see Methods for details). The mean spike density plots are shown for 1) all trials (bottom trace) 2) top 10 % activity corresponding to each time step (top trace) and 3) top 10 % activity from 80-180 ms aligned to the landmark-shift (middle trace). Note (the red histograms below the spike density plots) that the delay period possessed substantial spatially tuned neural activity along the T-T’ continuum, approximately 50 % of the neurons were tuned. A similar trend was noticed along the T-G continuum (not shown). **Figure 11B** shows the data (mean ± SEM) in the corresponding time steps for the population along the T-G (blue) and the T-T’ (red) continua. The solid circles indicate a significant shift from T (p < 0.005, one sampled Wilcoxon signed rank test), whereas the empty circle indicates a non-significant shift. The T-G code showed a significant shift at all the steps as reported previously (Bharmauria et al., 2020; Sajad et al., 2015). The T-T’ fits were slightly shifted from T at the first step, but this shift was significantly embedded only at the 4^th^ step (p = 0.02), then it shifted back at the 5^th^ step before significantly shifting toward T’ at the 6^th^ (p = 0.001) and 7^th^ (p = 0.003) steps when the gaze was just imminent.

To further tease apart the contribution of different cell types to the embedding of landmark influence, we divided the population into visual only (V), visuomotor (VM) and motor (M) cell types. The V neurons (n = 6, not shown) did not display any significant shift at any of the time steps, therefore they were eliminated from further analyses. **Figure 11C** shows the spike density plots as shown for the population in **Fig. 11A**. We did not notice any significant shift in the delay period along the T-T’ continuum for the VM neurons **(Figure 11D).** For the delay activity **(Fig. 11E)** in M neurons, along the T-T’ continuum, a significant shift was observed at the 3^rd^ step (p = 0.04), then the code shifted back before significantly shifting toward T’ at the 7^th^ step (p = 0.02) with impending gaze.

### Integration of the allocentric shift with the egocentric code

Until this point we observed that the landmark shift influences the motor code along the T-T’ coordinates, but we still need to address if there is any relation between the T-G and T-T’ transformations. To address this, post landmark-shift until the perisaccadic burst, we plotted the T-T’ score as a function of the corresponding T-G score for each neuron that exhibited spatial tuning for both **(Fig. 12A**). As for FEF (Bharmauria et al., 2020). we made following predictions for the embedding of the allocentric influence with the egocentric coordinates **(Fig. 12B):** 1) no influence, i.e., the coding was purely egocentric but as we have shown above that is not the case, 2) independent, the egocentric and allocentric codes are completely independent of each other, 3) fully integrated, the allocentric influence varies as a function of G, and 4) partial integration, a mix of 2 and 3.

**Figure 12.**
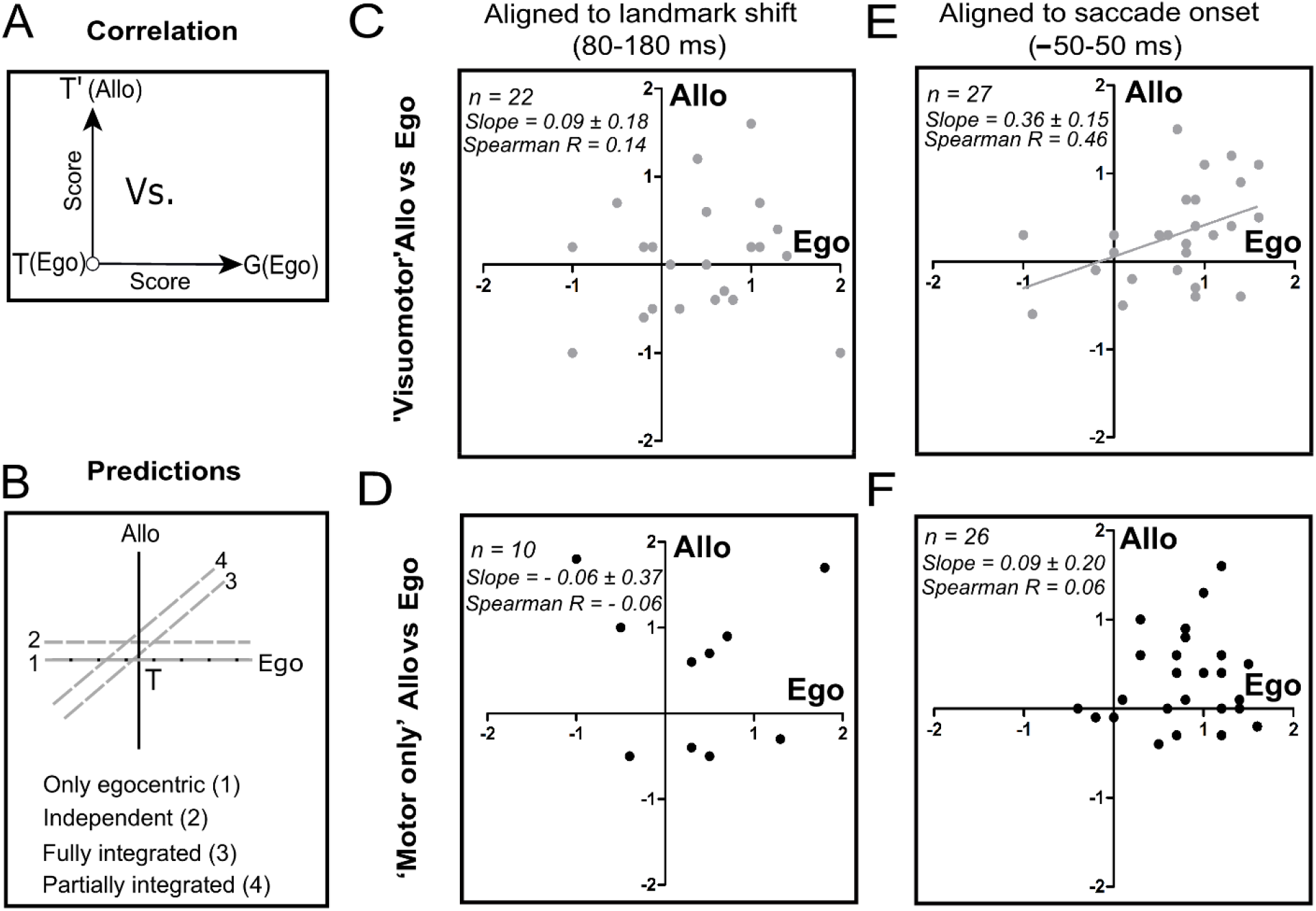
Correlation between T-G (egocentric) and T-T’ (allocentric) scores. **(A)** Schematic drawing of T-T’ plotting as a function of T-G scores. **(B)** Predictions for the embedding of allocentric shift with the egocentric codes: 1) only egocentric, 2) independent, 3) fully integrated, or 4) partially integrated. **(C)** No significant correlation between the corresponding T-T’ and T-G scores in the early post landmark-shift response for the VM (n = 22, Spearman R = 0.14, slope = 0.09 ± 0.18, p = 0.60) and (**D)** M neurons (n = 10, Spearman R = − 0.06, slope = −0.06 ± 0.37, p = 0.87). **(E)** A significant correlation between the corresponding T-T’ and T-G scores of VM neurons in the peri-saccadic burst (n = 27, Spearman R = 0.46, slope = 0.36 ± 0.15, p = 0.02). (F) No significant correlation in the perisaccadic burst between the T-T’ and T-G scores for M neurons (n = 26, Spearman R = 0.06, slope = 0.09 ± 0.20, p = 0.64).

We first did this analysis for early post-shift visual response in the 80-180 ms window of analysis for the VM and M neurons **(Fig. 12C-D).** No significant correlation was noticed for both the VM (**Fig. 12C;** Spearman R = 0.14; Slope = 0.09 ± 0.18, intercept = 0.03 ± 0.16, p = 0.60) and the M (**Fig. 12D;** Spearman R = − 0.06; Slope = − 0.06 ± 0.37, intercept = 0.52 ± 0.32, p = 0.87) neurons suggesting no integration in this period. We further plotted the correlation for the delay activity from early post-shift response until the saccade onset (roughly corresponding to steps 2-7 from the previous figure) for the whole population (M + VM) and the individual M and VM populations. We found no significant correlation at each of these steps either for the entire population and the sub-populations, implying that yet integration had not occurred, although a shift was noticed in the delay and the impending saccadic activity of M neurons **(Fig. 11F).** Finally, a significant correlation between the T-T’ and T-G was noticed for the VM (**Fig. 12E**, n = 27; Spearman R = 0.46, Slope = 0.36 ± 0.15, intercept = 0.05 ± 0.13, p = 0.02) neurons in the perisaccadic burst (−50-50 ms), but not for the M (**Fig. 12E**, n = 26; Spearman R = 0.06, Slope = 0.09 ± 0.20, intercept = 0.27 ± 0.18, p = 0.64) neurons. After combining the M and the VM neurons, a significant correlation still existed (Spearman R = 0.28, Slope = 0.26 ± 0.12, intercept = 0.13 ± 0.11, p = 0.03).

### Comparison with Spatiotemporal Integration in FEF

As noted above, we performed SEF and FEF recordings concurrently, providing an opportunity to compare the current dataset with the FEF dataset published previously published (Bharmauria et al. 2020), focusing on the spatiotemporal progression of ego – allocentric integration after the landmark shift. To do this, we performed a 3-factor ANOVA analysis on the visuomotor and motor populations from both areas (F1 = FEF/SEF, F2 =M/VM, F3 = time step). We found a significant difference between the VM and M neurons along the T-G (p = 0.009), T-T’ (p = 0.04) continua. We found no significant difference between the SEF and FEF along the T-G continuum (p = 0.10, suggesting similar egocentric transformations), but we found a significant difference along the T-T’ continuum (p = 0.009), implying a difference in allocentric processing. Moreover, a significant interaction was also noticed between the VM/M neurons of FEF/SEF (p = 0.04) along the T-T’ continuum. Finally, the M neurons of SEF displayed a significant shift in their delay activity (4^th^ step, Bonferroni corrected Mann-Whitney U-test, p = 0.006) towards T’ compared with the M neurons of FEF. These statistics support the observation that both areas showed transient T-T’ shifts in delay activity, but this primarily occurred in visuomotor neurons in the FEF (Bharmauria et al., 2020), as opposed to motor neurons in SEF. Finally, we note that whereas only SEF visuomotor neurons showed T-G / T-T’ correlation during the saccade burst **(Figure 12),** both visuomotor and motor neurons showed this correlation in the FEF (Bharmauria et al., 2020).

## DISCUSSION

This study addressed a fundamental question in cognitive neuroscience: how does the brain represent and integrate allocentric and egocentric spatial information? We used a cue-conflict memory-guided saccade task, in which a visual landmark shifted after a mask, to establish the basic egocentric coding mechanisms used by the SEF during head-unrestrained gaze shifts, and investigate how allocentric information is incorporated into these transformations. We found that: 1) Despite the presence of a visual landmark, spatially tuned SEF neurons predominantly show the same eye-centered codes as the FEF and SC (Sadeh et al., 2015; Sajad et al., 2015), i.e. target coding (T) in the visual burst and gaze position coding (G) in the motor burst. 2) After the landmark shift, motor neuron delay activity showed a transient shift in the same direction (T-T’). 3) A second perisaccadic shift was observed in visuomotor neurons. 4) Only the latter shift was correlated with T-G. Overall, the SEF showed similar egocentric visuomotor transformations, however, it integrated the landmark information into this transformation in a manner complementary to the FEF. Briefly, the novel results of this investigation implicate the SEF (and thus the frontal cortex) in the integration of allocentric and egocentric visual cues.

### General SEF Function: Spatial or Non-Spatial?

The FEF, LIP and SC show (primarily) contralateral visual and motor response fields involved in various spatial functions for gaze control (Andersen et al., 1985; Munoz, 2002; Schall, 1991; Schlag and Schlag-Rey, 1987), but the role of SEF is less clear (Purcell et al., 2012; Abzug and Sommer, 2018). Only 27% of our SEF neurons were spatially tuned, lower than our FEF recordings (50%) in the same sessions (Bharmauria et al., 2020). This is consistent with previous studies (Purcell et al., 2012; Schall, 1991) and the notion that the SEF also has non-spatial functions, such as, learning (Chen and Wise, 1995) prediction error encoding (Amador et al., 2000; Schlag-Rey et al., 1997; So and Stuphorn, 2012), performance monitoring (Sajad et al., 2019) and decision making (Abzug and Sommer, 2018). The general consensus is that SEF subserves various cognitive functions (Stuphorn et al., 2000; Tremblay et al., 2002) while also representing multiple spatial frames (Abzug and Sommer, 2017; Martinez-Trujillo et al., 2004; Stuphorn, 2015). It should be noted that these diverse signals (Abzug and Sommer, 2018; Sajad et al., 2019; So and Stuphorn, 2012) may be prominent in many of the spatially untuned neurons that were rejected in our analysis. However, the possibility of these signals influencing the spatially tuned response fields cannot be eliminated here (Abzug and Sommer, 2017; Purcell et al., 2012; Sajad et al., 2019). It can be reasonably hypothesized that spatially tuned neurons integrate non-spatial signals from untuned neurons (Pruszynski and Zylberberg, 2019) with spatial signals and forward these integrated signals to FEF neurons, thereby influencing gaze behavior in real space, thus providing a mechanism for SEF to implement executive function as behavior.

### Egocentric Transformations in the Gaze System

In the gaze control system, the consensus is that eye-centered visual and motor codes predominate (Goldberg et al., 2002; Klier et al., 2001; Paré and Wurtz, 2001; Russo and Bruce, 1993; Tehovnik et al., 2000), but alternative views persist (Caruso et al., 2018; Mueller and Fiehler, 2017). Visual-motor dissociation tasks (e.g., antisaccades) found that visual and motor activities coded target and saccade direction, respectively (Everling and Munoz, 2000; Sato and Schall, 2003; Takeda and Funahashi, 2004). However, this requires additional training and signals that would not be present during ordinary visually guided saccades (Amemori and Sawaguchi, 2006; Medendorp et al., 2005; Munoz and Everling, 2004), and the head was fixed in most such studies. We have previously extended these results to natural head-unrestrained gaze shifts in the SC and FEF (Sadeh et al., 2015; Sajad et al., 2015: Sadeh et al., 2020) and here in the SEF.

Consistent with previous reports we found that SEF response fields are primarily organized in eye-centered coordinates (Park et al., 2006; Russo and Bruce, 1993), and participate in progressive Target-to-Gaze transition like FEF and SEF (Sadeh et al., 2020; Sajad et al., 2016). Note that in the current study, deviations of gaze from the target (used to fit response fields against G) were produced in part by the landmark shift. However, this alone does not likely explain the T-G transition in our cells, because it happened continuously through the task, it was spatially separable and often uncorrelated with the neural response to the landmark shift (discussed below), and much of the gaze errors used to calculate this transition were not due to the landmark shift, but appeared to be due to general internal ‘noise’, as in our previous studies (Sadeh et al., 2020; Sajad et al., 2020, 2016) which was even larger without a landmark (Li et al., 2017). In short, the landmark shift clearly contributed to gaze errors, but cannot alone explain the T-G transition we observed here in SEF cells.

Besides the general resemblance of the T-G transition in SC, FEF, and SEF, we were able to compare the latter two directly in the current experiments, and found no significant difference. This level of spatial simplicity and homogeneity across these gaze-related areas likely results from shared inputs and extensive interconnectivity, and perhaps serves as a common baseline signal to carry more subtle cognitive modulations (Munoz, 2002; Munoz and Everling, 2004; Schall, 2015). This does not preclude different influences on spatial behavior, because 1) these structures may carry other more subtle spatial signals that might be decoded downstream (Bulkin and Groh, 2006; Gandhi and Katnani, 2011) and 2) the influence of these signals on behavior depends on how they project to downstream motor structures, and through further modulations such as ‘gain fields’ (Andersen and Buneo, 2002; Blohm and Crawford, 2009; Smith and Crawford, 2005). For example, microstimulation of the SC induces eye-fixed gaze shifts (Gandhi and Katnani, 2011; Klier et al., 2001; Wurtz and Albano, 1980) whereas microstimulation of frontal cortex produces gaze shifts toward a spectrum of eye, head, and body-centered goals (Abzug and Sommer, 2017; Martinez-Trujillo et al., 2004; Monteon et al., 2013; Sato and Schall, 2003; Schlag and Schlag-Rey, 1987) suggesting more complex motor transformations.

### Integration of Landmark-Centered Codes with Egocentric codes

Previous humans studies suggest that egocentric and allocentric codes are initially separated in the visual system (Chen et al., 2018, 2014; Milner and Goodale, 2006; Schenk, 2006) and then reintegrated in parietofrontal cortex (Chen et al., 2018). Relatively few neurophysiological studies have aimed at discriminating egocentric/allocentric codes in visual and visuomotor systems (Dean and Platt, 2006; Uchimura, M; Kumano, H; Kitazawa, 2017). Our recent study confirmed the role of the frontal eye fields in integrating these codes for gaze (Bharmauria et al., 2020). Specifically, the landmark shift paradigm used here induced an initial transient T-T’ shift in delay activity, followed by a shift that was integrated with the egocentric T-G code in the motor burst.

In some ways our SEF and FEF findings were similar, i.e., we found an influence of the landmark shift on the more basic egocentric codes, first as a multiplexed but independent code, and finally as an integrated influence within the egocentric motor code. However, there were mechanistic differences between these structures: the landmark shift was coded initially in SEF motor neurons (vs. visuomotor neurons in the FEF) and integration only occurred in the visuomotor burst (vs. all motor responses in FEF) suggesting a complementary mechanism between these areas.

In both the FEF and SEF, there was good agreement between the degree of landmark influence on the neural signals (measured either as a T-T’ shift or the final T-T’/T-G slope) and the actual gaze behavior, suggesting that these structures participate in the optimal integration described in previous studies, where ‘optimal’ is defined as the best estimate of target direction based on statistical weighing of signal uncertainty (Byrne and Crawford, 2010; Karimpur et al., 2020; Körding and Wolpert, 2004). This is consistent with the general notion that the brain employs Bayesian methods of statistical learning, using probabilistic strategy from the task/target distribution and feedback uncertainty, to optimize performance (Aagten-Murphy and Bays, 2019; Byrne and Crawford, 2010; Karimpur et al., 2020; Körding and Wolpert, 2004; Mutluturk and Boduroglu, 2014).

Finally, at first glance our findings contradict the classic findings of object-centered coding in the SEF (Olson and Gettner, 1995; Tremblay et al., 2002). However, there are important differences: besides being head-restrained, those studies involved explicit training and coding of one part of an object relative to other part, whereas, ours involved untrained and implicit coding of a target relative to a background landmark. Taken together, the SEF plays a role in both the implicit and explicit use of allocentric cues for gaze coding.

### A circuit model for allocentric/egocentric integration

**Fig. 13** provides a hypothetical circuit model for *allocentric / egocentric integration* in frontal cortex **(Fig. 13)**, based on our current results, FEF recordings obtained in the same recording sessions (Bharmauria et al., 2020), and previous literature. First, our data show that the FEF and SEF are initially driven by low latency (~80ms) egocentric visual inputs (target-in-eye coordinates), whereas the landmark influence has a higher latency (~200-300ms), consistent with more complex visual processing pathway. Such signals are thought to arise in the ventral visual stream (Chen et al., 2014; Milner and Goodale, 2006; Schenk, 2006) which projects to parietal cortex (Milner, 2017; Budisavljevic et al., 2018). Consistent with this, saccade-related activity in human parietal cortex is modulated by landmarks (Chen et al. 2017). Direct inputs to the prefrontal gaze system include LIP (Andersen et al., 1990; Schall et al., 1993; Stuphorn, 2015), and nuclei in central thalamus that relay inputs from the SC and substantia nigra pars reticulata (SnPR) (Lynch et al., 1994; Parent and Hazrati, 1995).

**Figure 13.**
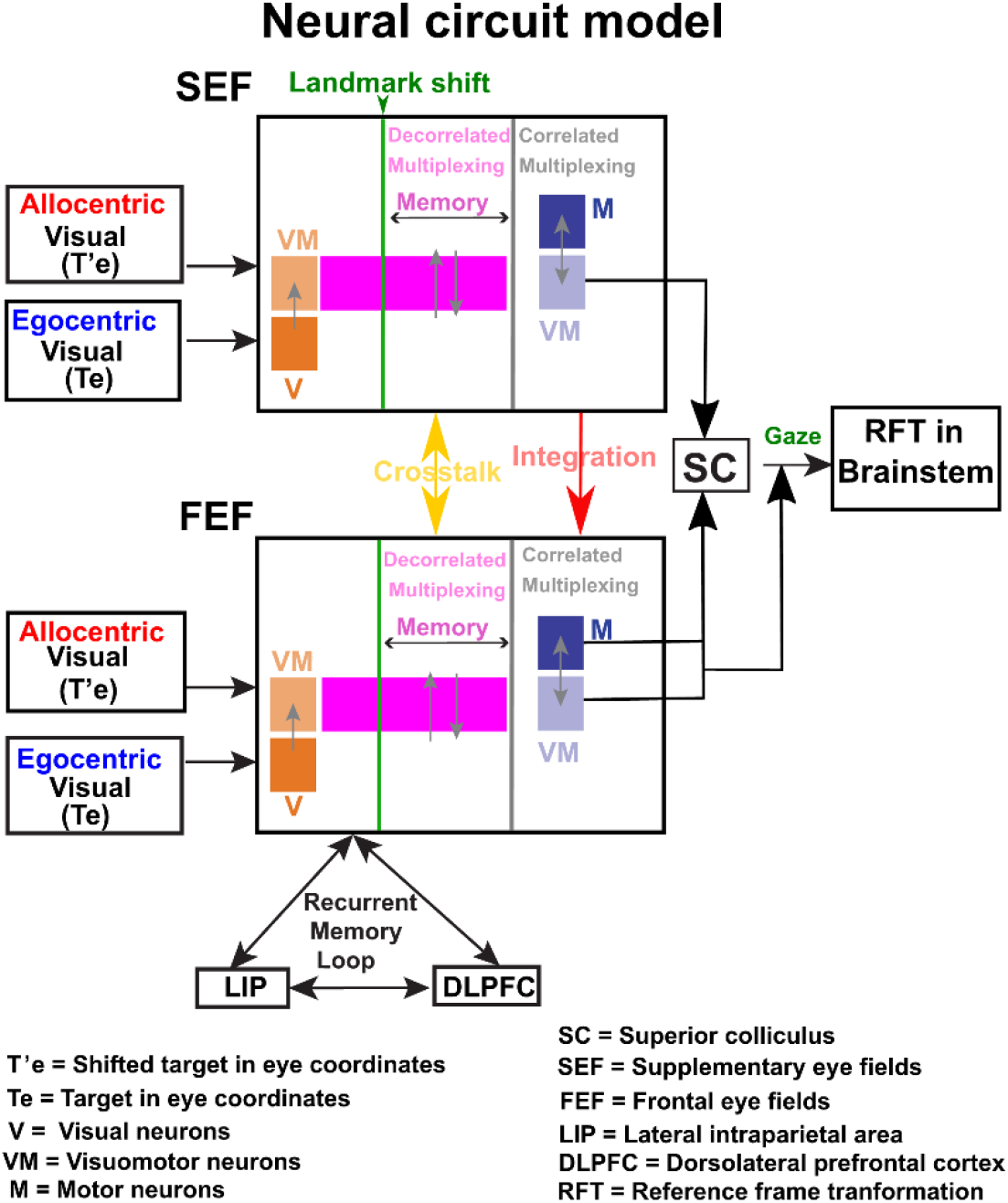
A neural circuit model for egocentric/allocentric multiplexing of signals in the FEF and SEF. The egocentric (Te) and allocentric information (T’e) arrive through the dorsal and ventral streams respectively and enter the FEF and the SEF as two separate visual inputs —egocentric (Te) input to the visual (V, dark orange) neurons and allocentric input to visuomotor neurons (VM, light orange). This processed visual information through the fronto-striato-thalamic loop (Chatham and Badre, 2015) is then relayed to the memory network (purple) with a considerable crosstalk (yellow arrow) between the FEF and SEF visuomotor-motor memory circuit, wherein the decorrelated scaling/multiplexing of the landmark-shift signal occurs with the egocentric flow. Finally, the first integrated signal arrives at the FEF motor circuitry from the most recent multiplexed memory (through continuous coordination between visuomotor-motor memory circuits of the FEF and the SEF), becoming fully integrated in the final motor burst (gaze command) of both areas (notably only SEF visuomotor neurons contribute). The final outputs project to SC and directly to the brainstem, which implements further reference frame transformations (RFT) from visual to motor coordinates for eye and head motion. **Note:** As the LIP, DLPFC and FEF share working memory loops, these brain areas may continuously provide egocentric/allocentric multiplexing signals. The gray arrows indicate the information flow between neuronal classes.

Second, we adopt the general convention that the SEF is involved more in executive control, whereas the FEF is more closely linked with eye control (Abzug and Sommer, 2018, 2017; Sajad et al., 2019; Stuphorn, 2015; Stuphorn et al., 2010). As discussed above, the non-spatial aspects could be subserved by a progression from non-spatial SEF neurons to spatial SEF and hence FEF neurons. This directional flow seems to hold for ego / allocentric integration, because in our data integration was more complete in the FEF motor burst than in the SEF motor burst. In this context, an executive control mechanism might explain why different contexts influence allocentric weighting (Neggers et al., 2005; Byrne and Crawford, 2010; Fiehler et al., 2014; Klinghammer et al., 2017)

Based on the preceding assumptions, we speculate that the SEF provides control signals to the FEF for context-appropriate reference frame integration. In this scheme, the landmark-shift is first assessed in SEF preparatory activity **(Fig. 11F)** and then relayed (red arrow) to FEF delay activity (Bharmauria et al., 2020). In the FEF, the egocentric-allocentric conflict was multiplexed, but not integrated (no correlation between T-G and T-T’) in the earlier delay activity of visuomotor neurons, however, an integration was observed in both the visuomotor and motor neurons in the final motor burst (Bharmauria et al., 2020). On the contrary, in the SEF, the multiplexed (non-integrated) signal first appeared in the delay activity of motor neurons and later (motor burst) integration only occurs in the visuomotor activity. Thus, in both areas, allocentric and egocentric signals are initially multiplexed in a decorrelated state within the reciprocal FEF-SEF loop (yellow arrows) but perhaps other working memory circuits are also involved (Christophel et al., 2017; Pinotsis et al., 2019). Finally, these signals become integrated in a path relayed from SEF visuomotor neurons to a more complete integration in the FEF motor response (red arrow), consistent with the latter being closer to motor output (Isa and Sasaki, 2002). We also speculate that the FEF and SEF neuronal classes have opposing roles with strongly and closely wired memory-related inter-class circuitries for a final gaze command. Briefly, we speculate that the SEF influences the final FEF burst in a learned, task-related capacity, although we cannot show that here. Thus the SEF relatively has more an ‘executive’ control over the ‘motor’ role of the FEF for goal-directed behavior.

This speculative model makes specific predictions for SEF-FEF spike correlation, i.e., 1) spatially tuned neurons should correlate across structures, 2) output neurons would correlate best with work-related activity in the opposite structure, 3) during the delay, SEF motor neurons should correlate best with FEF visuomotor neurons, 4) during saccades, SEF visuomotor neurons would correlate best with the FEF motor burst, and 5) neuron pairs that show temporal spike correlations will also show correlated spatial codes along the T-G/T-T’ continua. Since approximately 80% of our SEF neurons were recorded in conjunction with FEF neurons during our experiments, this is a feasible goal for a future study.

### General implications and conclusion

Previous studies have suggested that egocentric and allocentric visual cues are optimally integrated for goal directed action (Byrne and Crawford, 2010; Karimpur et al., 2020). Here, we emulated this behavior in the gaze system, and found that the SEF (like the FEF) is involved in an eye-centered transformation of target signals into gaze signals while incorporating landmark-centered information (Bharmauria et al., 2020). Taken together, these results suggest a neurophysiological model for optimal egocentric-allocentric integration in animals and humans. This is relevant for understanding normal function in daily (normal) and abnormal behavior, where brain damage preferentially affects egocentric vs. allocentric mechanisms (Milner and Goodale, 2006; Schenk, 2006). Knowledge of their integration circuits, combined with neuroplasticity, might provide access to preserved visual functions through targeted rehabilitation strategies.

## Acknowledgement

This project was supported by a Canadian Institutes for Health Research (CIHR) Grant and the Vision: Science to Applications (VISTA) Program, which is supported in part by the Canada first Research Excellence Fund. VB, XY, and HW were supported by CIHR and VISTA. AS was supported by an Ontario Graduate Scholarship. JDC is supported by the Canada Research Chair Program.

## CONTRIBUTION

VB did the experiments, analyzed the data and wrote the manuscript. AS contributed to data analysis. XY helped in the technical aspects of the recording data. HW performed the surgeries and helped in neural recordings. JDC conceived the study and contributed to data analysis, writing and editing of the manuscript.

---

**Figure 5-1:**
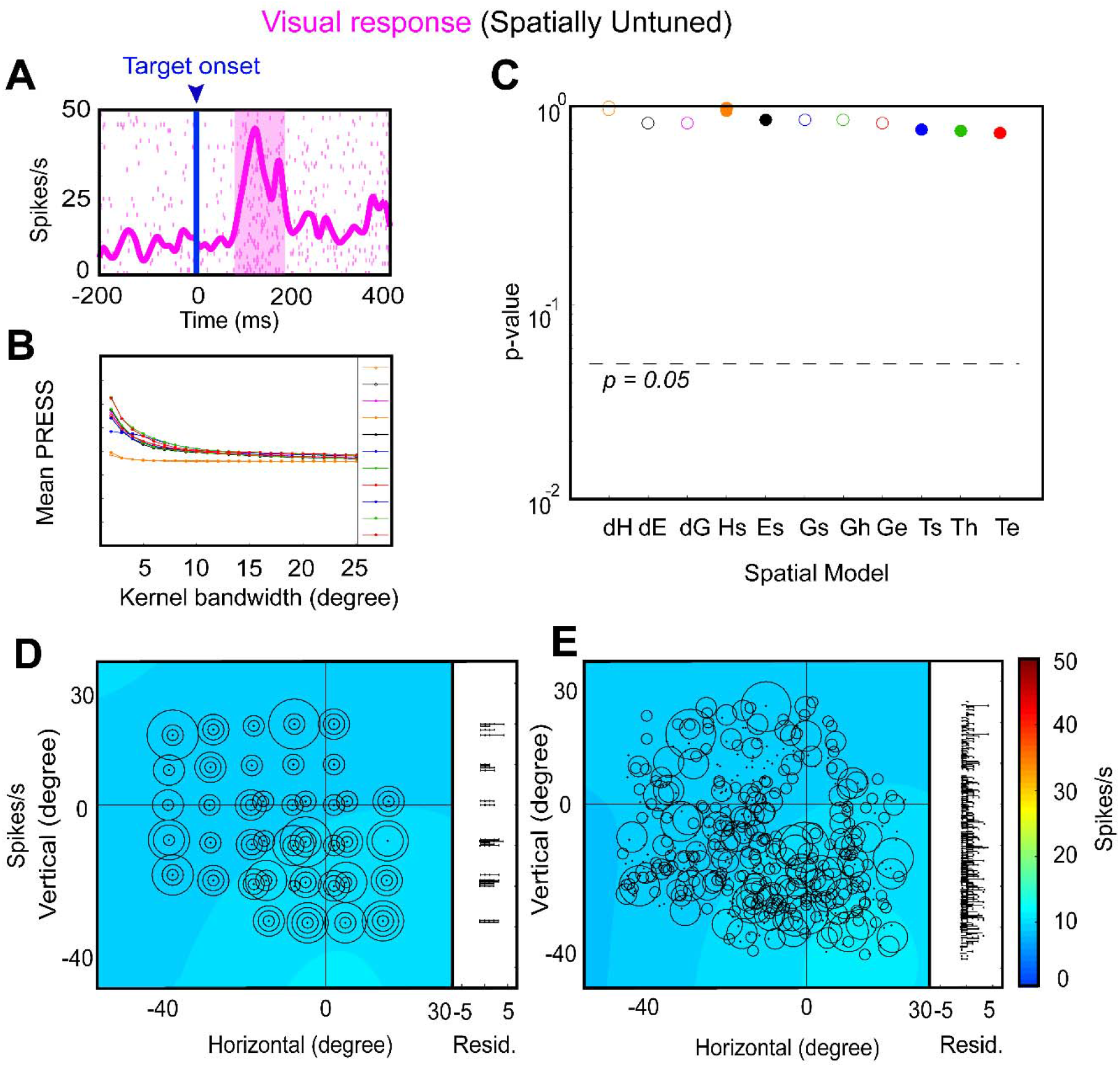
An example of a spatially untuned visual neuron. **(A)** Raster/spike density plot (with top 10 % responses) of the visual neuron aligned to the onset of target (blue arrow); the shaded pink region corresponds to the sampling window (80-180 ms) for response field analysis. **(B)** Mean residuals from the PRESS-statistics for all spatial models at different kernel bandwidths (2-25°). **(C)** The p-values statistics and comparison between different models. No model was significantly eliminated. **(D)** Representation of neural activity for Ts: target in space (screen). The circle corresponds to the magnitude of the response and heat map represents the non-parametric fit to these data. The corresponding residuals are displayed to the right **.(E)** Representation of the neural activity in Te (Target in eye).

**Figure 10-1:**
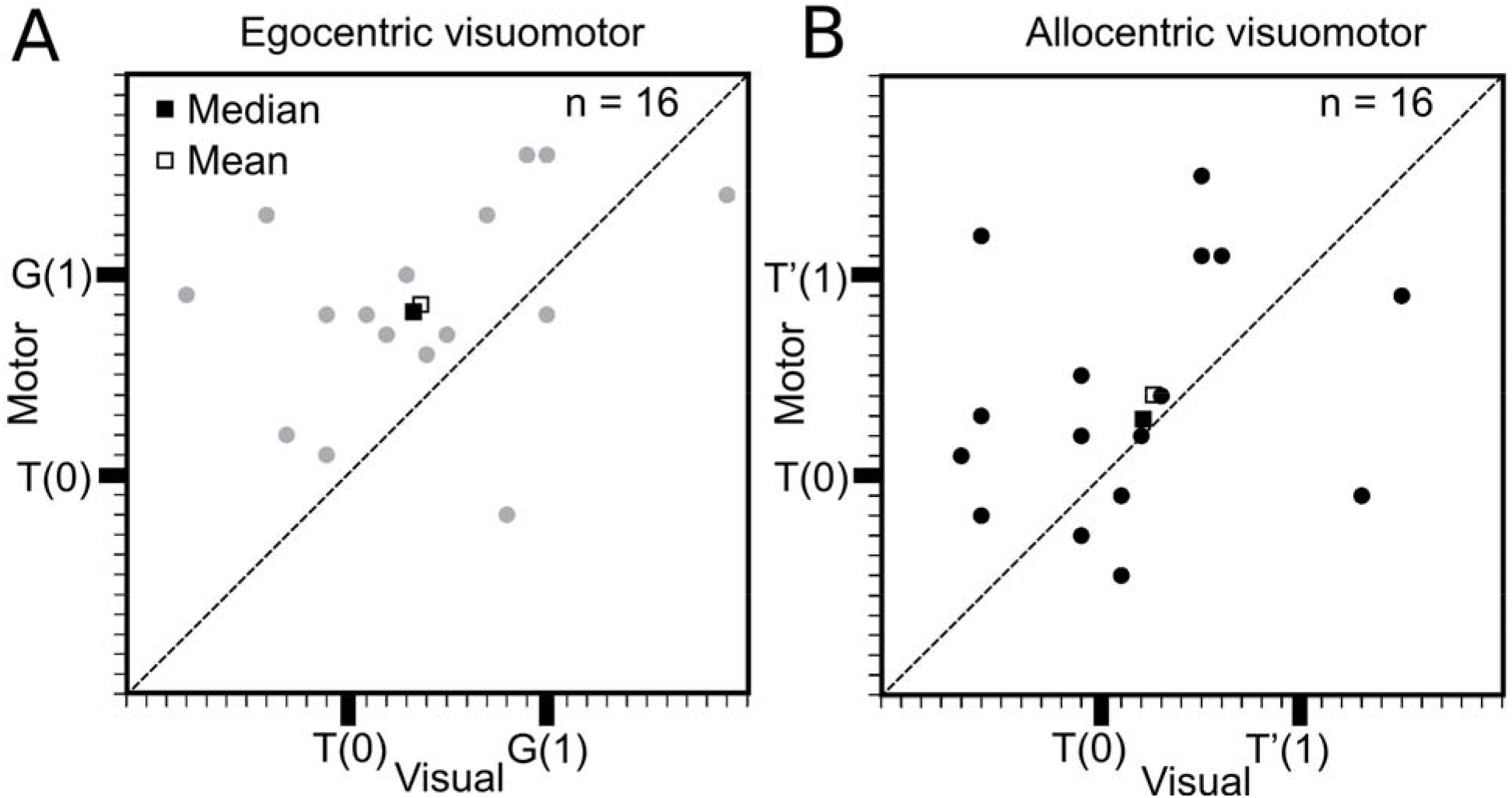
Visualmotor transformation at the single cell level. **(A)** Significant visual to motor transformation within the visuomotor neurons along the T-G continuum. **(B)** No significant visual to motor transformation along the TT’ continnum.

**Figure 11-1:**
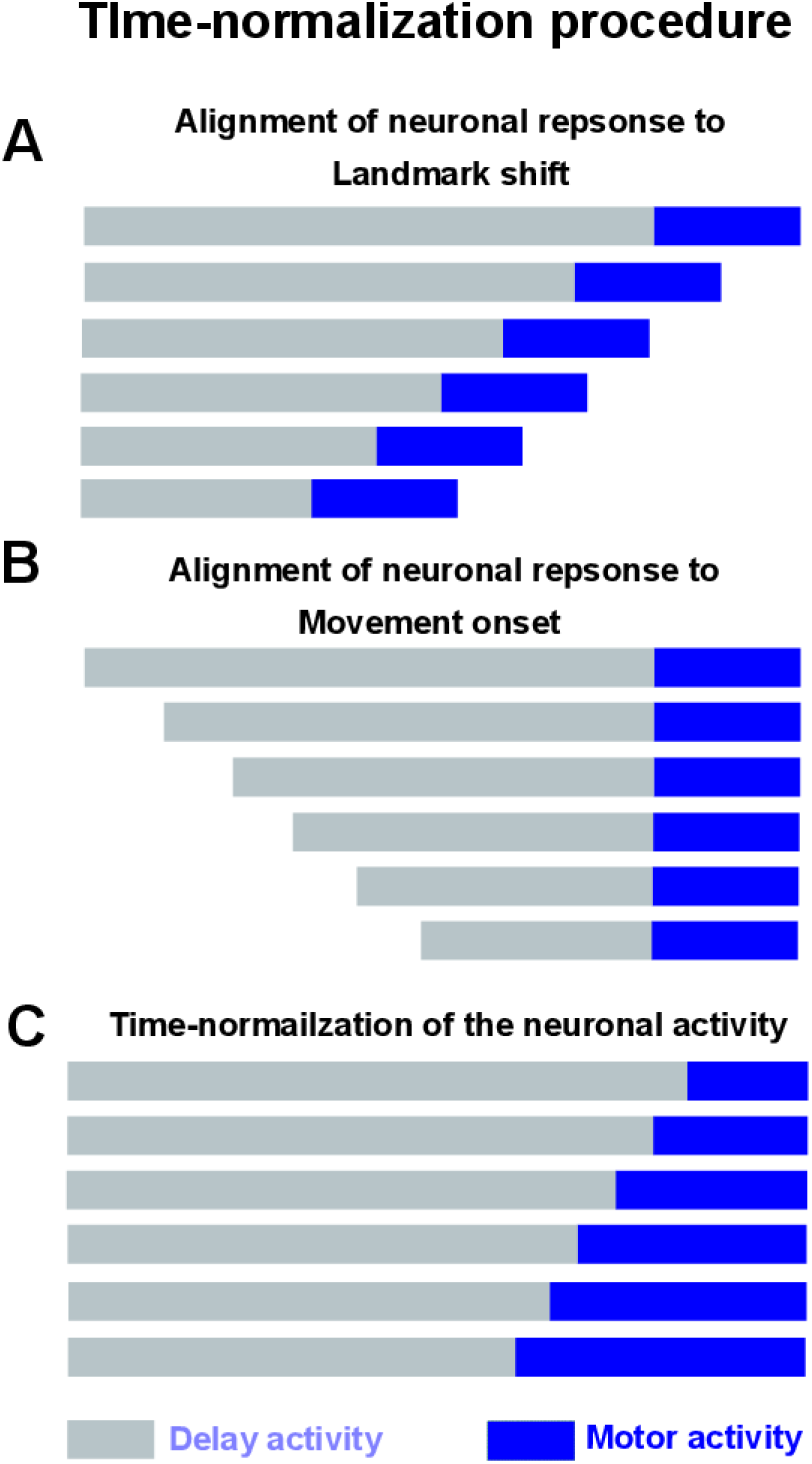
Time normalization procedure. **(A)** Alignment of responses to the landmark shift. **(B)** Alignment of responses to the saccade onset. Note: The alignment of responses in the standard was as in **A** and **B** leads to loss and/or mixing of responses thus not allowing us to track spatial codes through the entire trial across all trials. **(C)** Time-normalization from landmark shift until saccade onset. This procedure where we divide the activity into equal half-overlapping bins across all trials allows us to treat all the trials equally, thus as a continuum.

## Visual Abstract

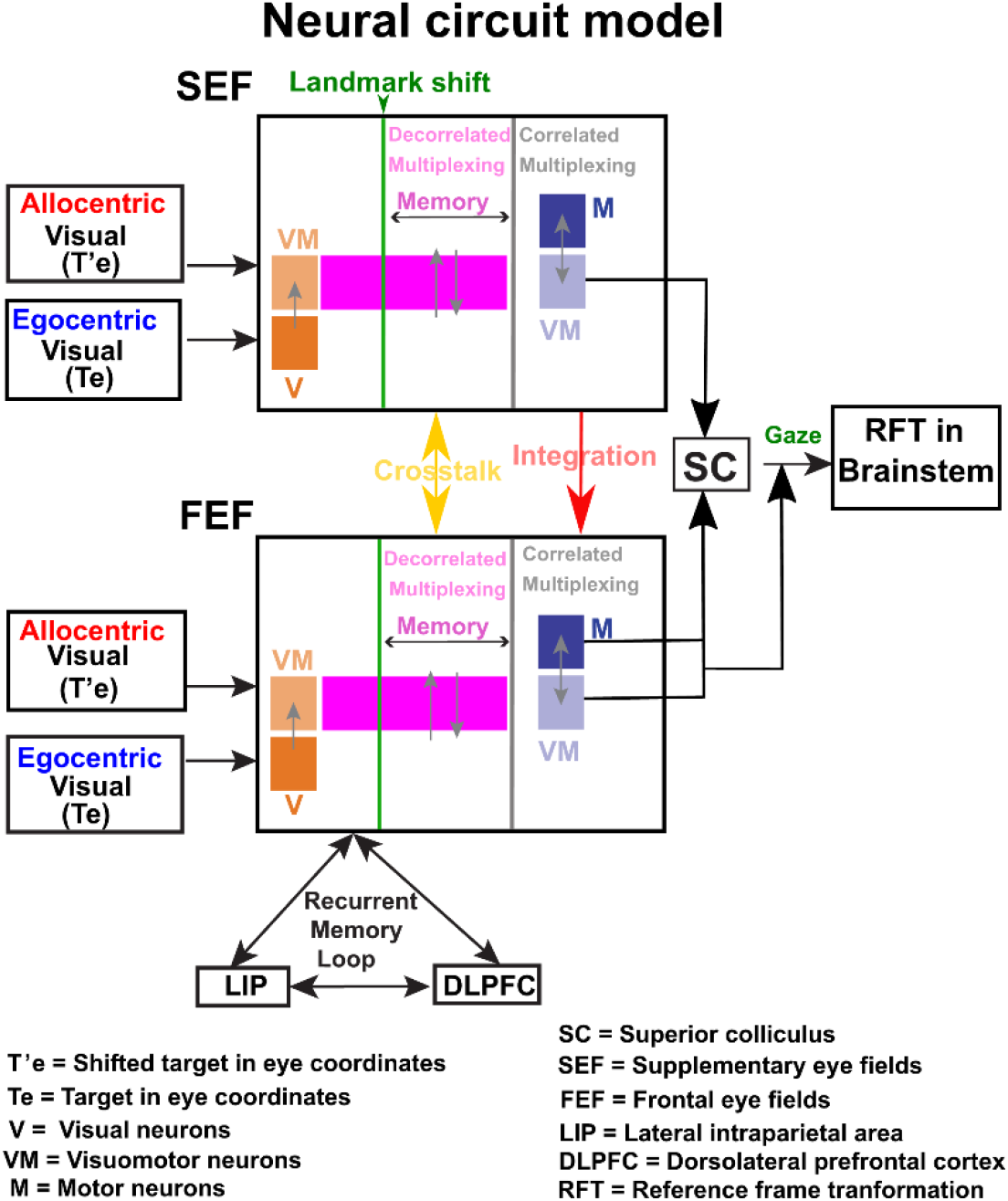

## References

Aagten-Murphy D, Bays PM (2019) Independent working memory resources for egocentric and allocentric spatial information. PLOS Comput Biol 15:e1006563.

Abzug ZM, Sommer MA (2018) Neuronal correlates of serial decision-making in the supplementary eye field. J Neurosci 38:7280–7292.

Abzug ZM, Sommer MA (2017) Supplementary Eye Fields[1] In: Reference Module in Neuroscience and Biobehavioral Psychology, Elsevier.

Amador N, Schlag-Rey M, Schlag J (2000) Reward-Predicting and Reward-Detecting Neuronal Activity in the Primate Supplementary Eye Field. J Neurophysiol 84:2166–2170.

Amemori KI, Sawaguchi T (2006) Rule-dependent shifting of sensorimotor representation in the primate prefrontal cortex. Eur J Neurosci 23:1895–1909.

Andersen RA, Buneo CA (2002) Intentional Maps in Posterior Parietal Cortex. Annu Rev Neurosci 25:189–220.

Andersen RA, Essick GK, Siegel RM (1985) Encoding of spatial location by posterior parietal neurons. Science (80-) 230:456–458.

Ball K, Smith D, Ellison A (2009) Both egocentric and allocentric cues support spatial priming in visual search. Neuropsychologia 47:1585–1591.

Bharmauria V, Bachatene L, Cattan S, Brodeur S, Chanauria N, Rouat J, Molotchnikoff S (2016) Network-selectivity and stimulus-discrimination in the primary visual cortex: Cell-assembly dynamics. Eur J Neurosci 43.

Bharmauria V, Sajad A, Li J, Yan X, Wang H, Crawford JD (2020) Integration of eye-centered and landmark-centered codes in frontal eye field gaze responses. Cereb Cortex bhaa090:https://doi.org/10.1093/cercor/bhaa090.

Blohm G, Crawford JD (2009) Fields of Gain in the Brain. Neuron 64:598–600.

Brandman DM, Cash SS, Hochberg LR (2017) Review: Human intracortical recording and neural decoding for brain computer interfaces. IEEE Trans Neural Syst Rehabil Eng 25:1687.

Bremmer F, Kaminiarz A, Klingenhoefer S, Churan J (2016) Decoding Target Distance and Saccade Amplitude from Population Activity in the Macaque Lateral Intraparietal Area (LIP). Front Integr Neurosci 10:30.

Bridgeman B, Peery S, Anand S (1997) Interaction of cognitive and sensorimotor maps of visual space. Percept Psychophys 59:456–469.

Bruce CJ, Goldberg ME (1985) Primate frontal eye fields. I. Single neurons discharging before saccades. J Neurophysiol 53:603–635.

Bruce CJ, Goldberg ME, Bushnell MC, Stanton GB (1985) Primate frontal eye fields. II. Physiological and anatomical correlates of electrically evoked eye movements. J Neurophysiol 54:714–34.

Bulkin DA, Groh JM (2006) Seeing sounds: visual and auditory interactions in the brain. Curr Opin Neurobiol.

Byrne PA, Crawford JD (2010) Cue Reliability and a Landmark Stability Heuristic Determine Relative Weighting Between Egocentric and Allocentric Visual Information in Memory-Guided Reach. J Neurophysiol 103:3054–3069.

Caruso VC, Pages DS, Sommer MA, Groh JM (2018) Beyond the labeled line: Variation in visual reference frames from intraparietal cortex to frontal eye fields and the superior colliculus. J Neurophysiol 119:1411–1421.

Chaplin TA, Hagan MA, Allitt BJ, Lui LL (2018) Neuronal Correlations in MT and MST Impair Population Decoding of Opposite Directions of Random Dot Motion. eNeuro 5.

Chatham CH, Badre D (2015) Multiple gates on working memory. Curr Opin Behav Sci 1:23–31.

Chen LL, Wise SP (1995) Neuronal activity in the supplementary eye field during acquisition of conditional oculomotor associations. J Neurophysiol 73:1101–1121.

Chen Y, Byrne P, Crawford JD (2011) Time course of allocentric decay, egocentric decay, and allocentric-to-egocentric conversion in memory-guided reach. Neuropsychologia 49:49–60.

Chen Y, Monaco S, Byrne P, Yan X, Henriques DYP, Crawford JD (2014) Allocentric versus egocentric representation of remembered reach targets in human cortex. J Neurosci 34:12515–26.

Chen Y, Monaco S, Crawford JD (2018) Neural substrates for allocentric-to-egocentric conversion of remembered reach targets in humans. Eur J Neurosci 47:901–917.

Christophel TB, Klink PC, Spitzer B, Roelfsema PR, Haynes J-D (2017) The Distributed Nature of Working Memory. Trends Cogn Sci 21:111–124.

Constantin AG, Wang H, Martinez-Trujillo JC, Crawford JD (2007) Frames of reference for gaze saccades evoked during stimulation of lateral intraparietal cortex. J Neurophysiol 98:696–709.

Crawford JD, Ceylan MZ, Klier EM, Guitton D (1999) Three-Dimensional Eye-Head Coordination During Gaze Saccades in the Primate. J Neurophysiol 81:1760–1782.

Crawford JD, Guitton D (1997) Visual-motor transformations required for accurate and kinematically correct saccades. J Neurophysiol 78:1447–67.

Dean HL, Platt ML (2006) Allocentric Spatial Referencing of Neuronal Activity in Macaque Posterior Cingulate Cortex. J Neurosci 26:1117–1127.

DeSouza JFX, Keith GP, Yan X, Blohm G, Wang H, Crawford JD (2011) Intrinsic reference frames of superior colliculus visuomotor receptive fields during head-unrestrained gaze shifts. J Neurosci 31:18313–26.

Ekstrom AD, Arnold AEGF, Iaria G (2014) A critical review of the allocentric spatial representation and its neural underpinnings: Toward a network-based perspective. Front Hum Neurosci 8:803.

Everling S, Dorris MC, Klein RM, Munoz DP (1999) Role of primate superior colliculus in preparation and execution of anti-saccades and pro-saccades. J Neurosci 19:2740–2754.

Everling S, Munoz DP (2000) Neuronal correlates for preparatory set associated with pro-saccades and anti-saccades in the primate frontal eye field. J Neurosci 20:387–400.

Fiehler K, Wolf C, Klinghammer M, Blohm G (2014) Integration of egocentric and allocentric information during memory-guided reaching to images of a natural environment. Front Hum Neurosci 8:636.

Gandhi NJ, Katnani HA (2011) Motor Functions of the Superior Colliculus. Annu Rev Neurosci 34:205–231.

Goldberg ME, Bisley J, Powell KD, Gottlieb J, Kusunoki M (2002) The role of the lateral intraparietal area of the monkey in the generation of saccades and visuospatial attention In: Annals of the New York Academy of Sciences, pp205–215. New York Academy of Sciences.

Goodale MA, Haffenden A (1998) Frames of reference for perception and action in the human visual system. Neurosci Biobehav Rev 22:161–72.

Goris RLT, Movshon JA, Simoncelli EP (2014) Partitioning neuronal variability. Nat Neurosci 17:858–865.

Huerta MF, Kaas JH (1990) Supplementary eye field as defined by intracortical microstimulation: Connections in macaques. J Comp Neurol 293:299–330.

Isa T, Sasaki S (2002) Brainstem control of head movements during orienting; organization of the premotor circuits. Prog Neurobiol 66:205–41.

Karimpur H, Kurz J, Fiehler K (2020) The role of perception and action on the use of allocentric information in a large-scale virtual environment. Exp Brain Res 1–14.

Keith GP, DeSouza JFX, Yan X, Wang H, Crawford JD (2009) A method for mapping response fields and determining intrinsic reference frames of single-unit activity: Applied to 3D head-unrestrained gaze shifts. J Neurosci Methods 180:171–184.

Khanna SB, Snyder AC, Smith MA (2019) Distinct Sources of Variability Affect Eye Movement Preparation. J Neurosci 39:4511–4526.

Klier EM, Wang H, Crawford JD (2003) Three-Dimensional Eye-Head Coordination Is Implemented Downstream From the Superior Colliculus. J Neurophysiol 89:2839–2853.

Klier EM, Wang H, Crawford JD (2001) The superior colliculus encodes gaze commands in retinal coordinates. Nat Neurosci 4:627–632.

Klinghammer M, Blohm G, Fiehler K (2017) Scene Configuration and Object Reliability Affect the Use of Allocentric Information for Memory-Guided Reaching. Front Neurosci 11:204.

Körding KP, Wolpert DM (2004) Bayesian integration in sensorimotor learning. Nature 427:244–247.

Leavitt ML, Pieper F, Sachs AJ, Martinez-Trujillo JC (2017) Correlated variability modifies working memory fidelity in primate prefrontal neuronal ensembles. Proc Natl Acad Sci U S A 114:E2494–E2503.

Lemay M, Bertram CP, Stelmach GE (2004) Pointing to an Allocentric and Egocentric Remembered Target in Younger and Older Adults. Exp Aging Res 30:391–406.

Li J, Sajad A, Marino R, Yan X, Sun S, Wang H, Crawford JD (2017) Effect of allocentric landmarks on primate gaze behavior in a cue conflict task. J Vis 17:20.

Martinez-Trujillo JC, Medendorp WP, Wang H, Crawford JD (2004) Frames of reference for eye-head gaze commands in primate supplementary eye fields. Neuron 44:1057–66.

Medendorp WP, Goltz HC, Vilis T (2005) Remapping the Remembered Target Location for Anti-Saccades in Human Posterior Parietal Cortex. J Neurophysiol 94:734–740.

Milner D, Goodale M (2006) The Visual Brain in Action. Oxford University Press.

Monteon JA, Wang H, Martinez-Trujillo J, Crawford JD (2013) Frames of reference for eye-head gaze shifts evoked during frontal eye field stimulation. Eur J Neurosci 37:1754–65.

Mueller S, Fiehler K (2017) Gaze-centered coding of proprioceptive reach targets after effector movement: Testing the impact of online information, time of movement, and target distance. PLoS One 12:e0180782.

Munoz DP (2002) Commentary: Saccadic eye movements: overview of neural circuitry. Prog Brain Res 140:89–96.

Munoz DP, Everling S (2004) Look away: The anti-saccade task and the voluntary control of eye movement. Nat Rev Neurosci.

Mutluturk A, Boduroglu A (2014) Effects of spatial configurations on the resolution of spatial working memory. Attention, Perception, Psychophys 76:2276–2285.

Neely KA, Tessmer A, Binsted G, Heath M (2008) Goal-directed reaching: movement strategies influence the weighting of allocentric and egocentric visual cues. Exp Brain Res 186:375–384.

Neggers SFW, Schölvinck ML, van der Lubbe RHJ (2005) Quantifying the interactions between allo- and egocentric representations of space. Acta Psychol (Amst) 118:25–45.

O’Keefe J (1976) Place units in the hippocampus of the freely moving rat. Exp Neurol 51:78–109.

O’Keefe J, Dostrovsky J (1971) The hippocampus as a spatial map. Preliminary evidence from unit activity in the freely-moving rat. Brain Res 34:171–175.

Olson CR, Gettner SN (1995) Object-centered direction selectivity in the macaque supplementary eye field. Science 269:985–8.

Paré M, Wurtz RH (2001) Progression in neuronal processing for saccadic eye movements from parietal cortex area LIP to superior colliculus. J Neurophysiol 85:2545–2562.

Park J, Schlag-Rey M, Schlag J (2006) Frames of Reference for Saccadic Command Tested By Saccade Collision in the Supplementary Eye Field. J Neurophysiol 95:159–170.

Pinotsis DA, Buschman TJ, Miller EK (2019) Working Memory Load Modulates Neuronal Coupling. Cereb Cortex 29:1670–1681.

Pruszynski JA, Zylberberg J (2019) The language of the brain: real-world neural population codes. Curr Opin Neurobiol 58:30–36.

Purcell BA, Weigand PK, Schall JD (2012) Supplementary eye field during visual search: Salience, cognitive control, and performance monitoring. J Neurosci 32:10273–10285.

Rosenbaum RS, Ziegler M, Winocur G, Grady CL, Moscovitch M (2004) “I have often walked down this street before”: fMRI Studies on the hippocampus and other structures during mental navigation of an old environment. Hippocampus 14:826–835.

Russo GS, Bruce CJ (1993) Effect of eye position within the orbit on electrically elicited saccadic eye movements: A comparison of the macaque monkey’s frontal and supplementary eye fields. J Neurophysiol 69:800–818.

Sadeh M, Sajad A, Wang H, Yan X, Crawford JD (2020) Timing determines tuning: A rapid spatial transformation in superior colliculus neurons during reactive gaze shifts. eNeuro 7.

Sadeh M, Sajad A, Wang H, Yan X, Crawford JD (2015) Spatial transformations between superior colliculus visual and motor response fields during head-unrestrained gaze shifts. Eur J Neurosci 42:2934–2951.

Sajad A, Godlove DC, Schall JD (2019) Cortical microcircuitry of performance monitoring. Nat Neurosci 22:265–274.

Sajad A, Sadeh M, Crawford JD (2020) Spatiotemporal transformations for gaze control. Physiol Rep.

Sajad A, Sadeh M, Keith GP, Yan X, Wang H, Crawford JD (2015) Visual-Motor Transformations Within Frontal Eye Fields During Head-Unrestrained Gaze Shifts in the Monkey. Cereb Cortex 25:3932–52.

Sajad A, Sadeh M, Yan X, Wang H, Crawford JD (2016) Transition from Target to Gaze. Coding in Primate Frontal Eye Field during Memory Delay and Memory-Motor Transformation. eNeuro 3.

Sato TR, Schall JD (2003) Effects of stimulus-response compatibility on neural selection in frontal eye field. Neuron 38:637–648.

Schall JD (2015) Visuomotor Functions in the Frontal Lobe. Annu Rev Vis Sci 1:469–498.

Schall JD (1991) Neuronal activity related to visually guided saccades in the frontal eye fields of rhesus monkeys: Comparison with supplementary eye fields. J Neurophysiol 66:559–579.

Schall JD, Hanes DP, Thompson KG, King DJ (1995) Saccade target selection in frontal eye field of macaque. I. Visual and premovement activation. J Neurosci 15:6905–18.

Schall JD, Morel A, Kaas JH (1993) Topography of supplementary eye field afferents to frontal eye field in macaque: Implications for mapping between saccade coordinate systems. Vis Neurosci 10:385–393.

Schenk T (2006) An allocentric rather than perceptual deficit in patient D.F. Nat Neurosci 9:1369–1370.

Schlag-Rey M, Amador N, Sanchez H, Schlag J (1997) Antisaccade performance predicted by neuronal activity in the supplementary eye field. Nature 390:398–401.

Schlag J, Schlag-Rey M (1987) Evidence for a supplementary eye field. J Neurophysiol 57:179–200.

Smith MA, Crawford JD (2005) Distributed Population Mechanism for the 3-D Oculomotor Reference Frame Transformation. J Neurophysiol 93:1742–1761.

So N, Stuphorn V (2012) Supplementary eye field encodes reward prediction error. J Neurosci 32:2950–2963.

Stuphorn V (2015) The role of supplementary eye field in goal-directed behavior. J Physiol Paris.

Stuphorn V, Brown JW, Schall JD (2010) Role of supplementary eye field in saccade initiation: Executive, not direct, control. J Neurophysiol 103:801–816.

Stuphorn V, Taylor TL, Schall JD (2000) Performance monitoring by the supplementary eye field. Nature 408:857–860.

Takeda K, Funahashi S (2004) Population Vector Analysis of Primate Prefrontal Activity during Spatial Working Memory. Cortex December 14:1328–1339.

Tehovnik EJ, Sommer MA, Chou IH, Slocum WM, Schiller PH (2000) Eye fields in the frontal lobes of primates. Brain Res Rev.

Tremblay L, Gettner SN, Olson CR (2002) Neurons With Object-Centered Spatial Selectivity in Macaque SEF: Do They Represent Locations or Rules? J Neurophysiol 87:333–350.

Uchimura, M; Kumano, H; Kitazawa S (2017) Rapid allocentric coding in the monkey precuneus In: Socety for Neuroscience, p589.24/ GG19.

Wurtz RH, Albano JE (1980) Visual-Motor Function of the Primate Superior Colliculus. Annu Rev Neurosci 3:189–226.

Zylberberg J (2018) The role of untuned neurons in sensory information coding. bioRxiv 134379.

